# Non-canonical hepatic androgen receptor mediates glucagon sensitivity in female mice through the PGC1α/ERRα/mitochondria axis

**DOI:** 10.1101/2023.11.02.565247

**Authors:** Jie Chen, Yuanyuan Wu, Wanyu Hao, Jia You, Lianfeng Wu

## Abstract

The hormone glucagon plays a crucial role in maintaining normal blood glucose levels and has recently been found to promote liver lipid catabolism in both clinical and basic research studies. However, the mechanisms by which glucagon signaling synchronizes glucose and lipid metabolism in the liver remain poorly understood. Here, we report that hepatic androgen receptor (AR) functions as a critical mediator of glucagon signaling, contributing to sexual dimorphism in hepatic glucagon sensitivity. The inhibition of AR, either through chemical or genetic means, impairs the ability of glucagon to stimulate gluconeogenesis and lipid catabolism in primary hepatocytes. Notably, female mouse hepatocytes express up to three times more AR than their male littermates. Consequently, they respond to glucagon to a much greater extent in terms of glucose production and lipid catabolism outcomes. Moreover, liver-specific AR knockout impairs the effect of glucagon in inducing blood glucose production, especially in female mice. Mechanistically, we demonstrate that hepatic AR drives a PGC1α/ERRα-mitochondria axis to promote lipid catabolism for liver glucose production in response to glucagon treatment. Together, our study sheds light on the role of hepatic AR in mediating glucagon activity in orchestrating glucose and lipid metabolism, particularly in female mice. These findings may help elucidate the factors responsible for sex differences in glucagon sensitivity and the development of fatty liver disease.

## Main

Glucagon is widely recognized as a counter-regulatory hormone to insulin, responsible for maintaining blood glucose levels through the stimulation of hepatic glucose production. In addition to its role in hepatic glucose regulation, glucagon has been found as a pivotal regulator of various metabolic processes, including lipid and energy metabolism^1^. However, the clinical investigation of glucagon receptor (GCGR) antagonists as a treatment for hyperglycemia in type 2 diabetes, such as LY2409021 and PF-06291874, is hindered due to their potential association with an elevated risk of developing non-alcoholic fatty liver disease (NAFLD)^2,3^. Conversely, studies have demonstrated that chronic perfusion of glucagon at a physiologic dose can reduce hepatic fat accumulation in mice, leading to improved insulin sensitivity and decreased blood glucose levels^4^. Additionally, research has shown that activation of the glucagon receptor not only synergizes with the beneficial effects of GLP-1 receptor agonists but also enhances their efficacy in treating metabolic disorders such as NAFLD and obesity^5–7^. These findings suggest that glucagon may exert a significant impact on liver function by promoting lipid mobilization and augmenting gluconeogenesis.

Blood glucagon levels increase as a compensatory response to impaired glucagon signaling pathways. This phenomenon has been observed in various contexts, such as when the *Gcgr* gene is knocked out in the liver, where it is highly expressed^8^, or when the downstream signaling molecule of GCGR, Gs alpha, is eliminated from the liver^9^. Patients diagnosed with NAFLD frequently exhibit hyperglucagonemia^10–12^, which indicates the existence of a “glucagon resistance” condition. Improving glucagon sensitivity is immensely important in the treatment of NAFLD. Moreover, these strategies have the potential to address the underlying mechanisms responsible for the beneficial synergistic effects observed with the simultaneous administration of GCGR agonists and GLP-1 receptor agonists in managing metabolic disorders^5–7^. However, the precise role of glucagon in regulating hepatic lipid catabolism remains incompletely understood.

The androgen receptor (AR) is a nuclear hormone receptor that binds to androgen hormones, such as testosterone and dihydrotestosterone. It is expressed in diverse tissues, including the testes, prostate gland, and liver, and plays an essential role in male sexual development and function^13^. Apart from its well-established involvement in sexual development and the pathophysiology of prostate cancer, recent evidence suggests that the AR also participates in other functions, such as hepatic metabolism and disorders. Notably, studies have indicated that the elimination of hepatic AR can protect female mice from liver steatosis induced by persistent hyperandrogenemia, a pathological condition characterized by excessive levels of male hormones^14,15^. Conversely, in male mice fed a high-fat diet, the knockout of hepatic AR has been observed to exacerbate liver steatosis^16^. These contradictory findings imply that the role of hepatic AR in liver steatosis may vary in different pathological conditions or different sexes. Therefore, there is still a lack of comprehensive understanding regarding the function of hepatic AR in various contexts, mainly due to limited knowledge about its role in normal physiology.

The PGC1α/ERRα complex plays a crucial role as a regulator of lipid catabolism and energy metabolism^17^. PGC1α acts as a co-activator of ERRα, significantly augmenting its transcriptional activity^18,19^. ERRα directly binds to the promoters of genes involved in fatty acid oxidation, the tricarboxylic acid (TCA) cycle, and the electron transport chain, thereby controlling the expression of these genes^17,20^. Specifically, the PGC1α/ERRα complex coordinates the expression of these metabolic genes in specific tissues or under certain conditions in response to elevated energy demands. For instance, exposure to cold stimuli leads to the upregulation of ERRα and PGC1α in mouse brown adipose tissue and skeletal muscle, resulting in the activation of mitochondrial oxidative phosphorylation and subsequent heat production^21,22^. As a result, ERRα-deficient mice struggle to maintain normal body temperature when exposed to cold due to impaired mitochondrial oxidative capacity^23,24^. Exercise also induces the upregulation of ERRα and PGC1α in both animal models and humans^25,26^. Moreover, fasting has been shown to increase the levels of PGC1α and ERRα in the liver^27^. PGC1α can bind to specific transcription factors, such as HNF4, FOXO1, and GR, to directly enhance the expression of genes involved in gluconeogenesis ^28,29^. During fasting, the activation of lipid catabolism by glucagon is necessary to meet the energy requirements for gluconeogenesis^4^. Nevertheless, the exact role of the PGC1α/ERRα complex in this process has not been fully elucidated.

In this study, we unexpectedly discovered that enzalutamide, a clinically prescribed AR inhibitor, could effectively inhibit glucagon-stimulated gluconeogenesis. Genetic disruptions of hepatic AR through knockdown or knockout approaches could also markedly impair glucagon-stimulated gluconeogenesis in both primary hepatocytes and mice. Furthermore, we demonstrated that AR played a crucial role in regulating glucagon-stimulated lipid catabolism. Notably, we found that female mouse hepatocytes expressed significantly higher levels of AR than male mouse hepatocytes, which may explain the greater response of these cells to glucagon in inducing gluconeogenesis and lipid catabolism. In summary, our findings not only unveil the important role of AR in the liver as a regulator of glucagon signaling, which may elucidate the sex disparity in hepatic glucagon responsiveness, but also provide a potential target for future strategies aimed at modulating glucagon sensitivity.

## Results

### An unexpected role of hepatic AR in mediating glucagon response

The antidiabetic drug metformin, which is prescribed worldwide, is believed to inhibit hepatic gluconeogenesis as its mechanism of action^30,31^. To identify compounds that may act more efficiently than metformin, we conducted a high-throughput screen of 1,124 FDA-approved drugs using our previously established *C. elegans* metformin reporter^32^ (Extended Data Fig. 1a). The screen yielded 96 candidate compounds that positively regulated the expression of CeACAD10 (Extended Data Fig. 1b, c and Supplementary Table 1). Among the top 6 candidates that markedly increased the CeACAD10 expression (Extended Data Fig. 1c, d), even at concentrations one thousand times lower than that of metformin, two of them had their direct targets expressed in the liver according to annotations in Human Protein Atlas and had been reported to possess metabolic activity. These two compounds were the AR inhibitor, enzalutamide^33,34^ and the selective potentiator of cystic fibrosis transmembrane conductance regulator, VX-770^35,36^. Therefore, we chose these two compounds for glucose production testing in mouse primary hepatocytes, with the addition of glucagon as a gluconeogenesis stimulator. Significantly, we found that enzalutamide, but not VX-770, suppressed glucagon-stimulated glucose production, in a manner similar to that of metformin (Extended Data Fig. 1e).

To validate the specificity of enzalutamide, we utilized flutamide, another commonly used AR inhibitor (Fig. 1a, b), in the gluconeogenesis test. Additionally, we employed an adenovirus-mediated shRNA knockdown approach to decrease the expression of *Ar* (Fig. 1c, d and Extended Data Fig. 2a, b). The results showed that these additional treatments also significantly impeded glucagon-stimulated glucose production and the expression of genes involved in gluconeogenesis, such as the phosphoenolpyruvate carboxykinase 1 (*Pck1*) and the glucose-6-phosphatase catalytic subunit (*G6pc*), similar to the effects elicited by enzalutamide (Fig. 1a-d). This consistent outcome implicates that AR may play an important role in modulating glucose homeostasis during glucagon stimulation. Subsequently, we investigated whether the AR inhibitor enzalutamide could replicate the regulatory effect of metformin on blood glucose control in mice. However, contrary to the acute administration of metformin in mice that led to the expected reduction in basal blood glucose levels, we did not observe the anticipated effect when mice were acutely treated with enzalutamide (Extended Data Fig. 3a, b). These results suggest that enzalutamide may not serve as a compound that replicates the effect of metformin in regulating blood glucose homeostasis in an organism.

**Fig. 1.**
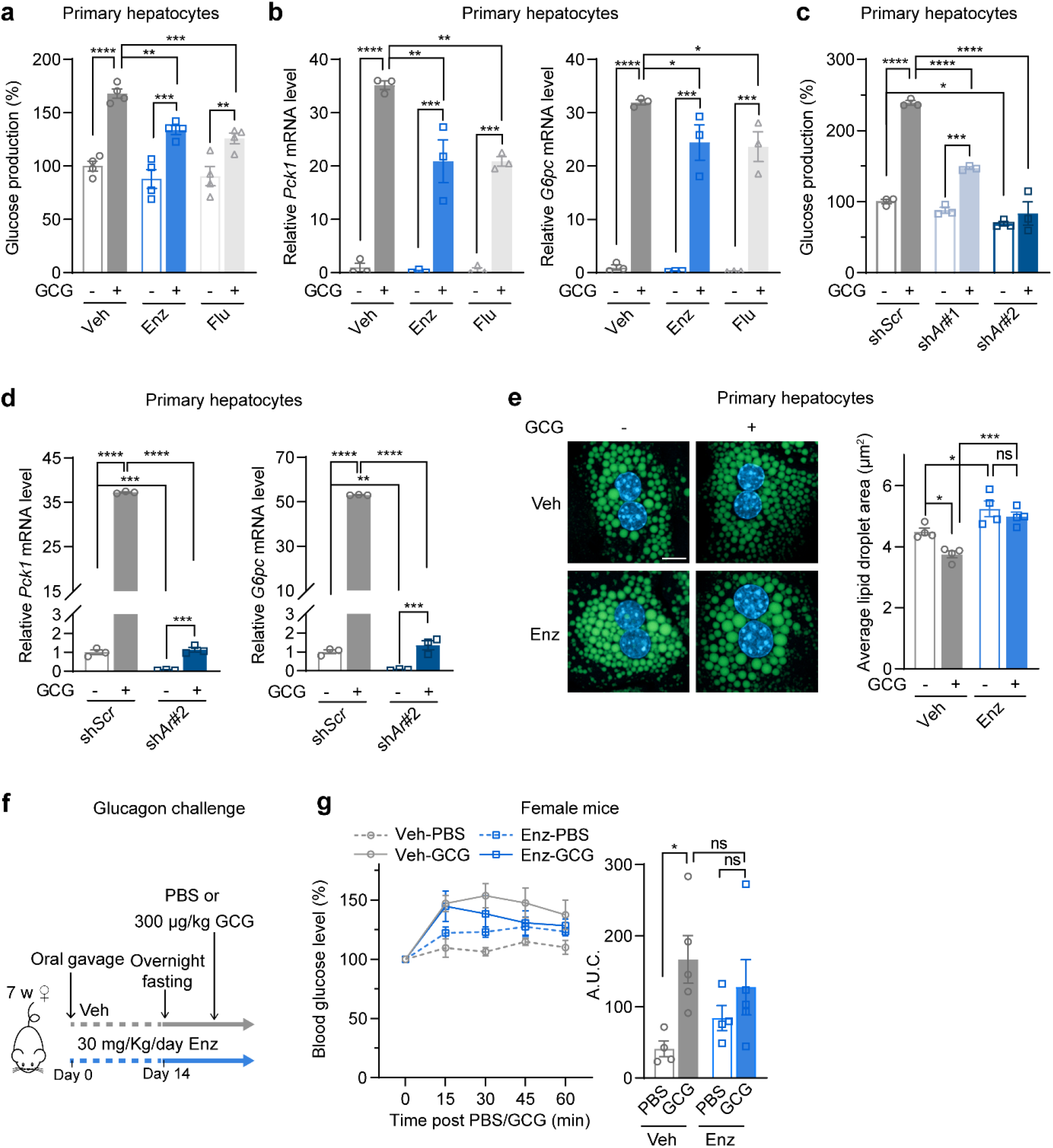
Inhibition of AR hampers the effects of glucagon on gluconeogenesis and lipid catabolism. **a, b,** Hepatic glucose production (**a**) and qRT-PCR analysis of gluconeogenic genes *Pck1* and *G6pc* (**b**) were conducted in female mouse primary hepatocytes pretreated with vehicle (Veh), 20 μM enzalutamide (Enz), or 20 μM flutamide (Flu) for 4 hours prior to treatment with either Veh or 20 nM glucagon (GCG) for 18 hours. **c, d,** Hepatic glucose production (**c**) and qRT-PCR analysis of gluconeogenic genes *Pck1* and *G6pc* (**d**) in primary hepatocytes isolated from female mice. The hepatocytes were infected with Ad-sh*Scr* or Ad-sh*Ar* and then stimulated with either Veh or 100 nM GCG for 6 hours. **e,** The hepatocyte lipid droplets were detected through BODIPY staining (left) and their sizes were quantified (right). This analysis was conducted using female mouse primary hepatocytes, which were pretreated with Veh or 30 μM Enz for 4 hours prior to treatment with Veh or 100 nM glucagon for 15 hours. Scale bars, 10 μm. A total of more than 3000 lipid droplets were evaluated per group, using 80-120 hepatocytes from 4 mice. **f, g,** Glucagon challenge assay was conducted using female mice that were administered either Veh or Enz via oral gavage for a duration of two weeks (n=4-5/group) (**f**). Blood glucose levels were measured before and after intraperitoneal administration of PBS or 300 μg/kg glucagon, at specified time points. The blood glucose of each mouse were normalized with basal blood glucose as 100%. The area under curve (A.U.C.) was then calculated adjacent to the glucose curve (**g**). The data were presented as mean ± SEM from three or more independent experiments or from mice with the indicated number. The statistical significance was determined using the following criteria: ****p≤0.0001, ***p≤0.001, **p≤0.01, *p≤0.05; ns denotes no significance with p > 0.05. The statistical tests employed were two-way ANOVA for (**a-e, g**).

A consistent and significant decrease in glucose production was observed in primary hepatocytes using different approaches to inhibit AR activity, under conditions of glucagon stimulation (Fig. 1a-d). This suggests that the inhibition of AR, either with the drug enzalutamide or through other means, may have disrupted glucagon’s activity in hepatocytes. Glucagon is known to play a major role in regulating various hepatic metabolic processes, including gluconeogenesis and lipid metabolism^1^. Hence, we investigated whether inhibiting AR could mitigate the effects of glucagon on fat accumulation in primary hepatocytes. As anticipated, treatment with enzalutamide not only resulted in an increase in lipid droplet sizes but also completely reversed the reductions in lipid droplet size induced by glucagon in primary hepatocytes from female mice (Fig. 1e). Recent studies have revealed that glucagon treatment affects both blood glucose levels and liver fat breakdown in mice^4^. To further explore this phenomenon, we administered enzalutamide to female mice via chronic oral gavage at a dosage of 30 mg/kg/day for 21 days. The mice tolerated this dosage well, as evidenced by their unaffected body weight growth (Fig. 1f and Extended Data Fig. 3c). Remarkably, enzalutamide administration almost entirely blocked the ability of glucagon to elevate blood glucose levels in female mice (Fig. 1g). Furthermore, the administration of enzalutamide in female mice resulted in a significant increase in liver fat accumulation (Extended Data Fig. 3d, e), which is consistent with the observed effects of glucagon signaling inhibition^37,38^. Collectively, these results demonstrate an unexpected and critical role of hepatic AR in mediating the response to glucagon.

### The higher expression of hepatic AR in female mice results in increased sensitivity to glucagon

In our assessment of AR expression in primary hepatocytes derived from male and female mice, we surprisingly found that both the mRNA and protein levels of AR were expressed at over three times higher levels in female mouse hepatocytes compared to their male littermates (Fig. 2a, b). Accordingly, greater glucagon responses were detected in female mouse primary hepatocytes compared to males. This was consistently demonstrated by the enhanced glucose production induced by glucagon (Fig. 2c), upregulation of gluconeogenesis genes (Fig. 2d), and reduction in lipid droplet sizes (Fig. 2e). Aligning with the expression pattern of AR in primary hepatocytes from both sexes, we observed significantly higher expression levels of AR at both the mRNA and protein levels in the liver tissues of female mice as compared to males (Fig. 2f, g).

**Fig. 2.**
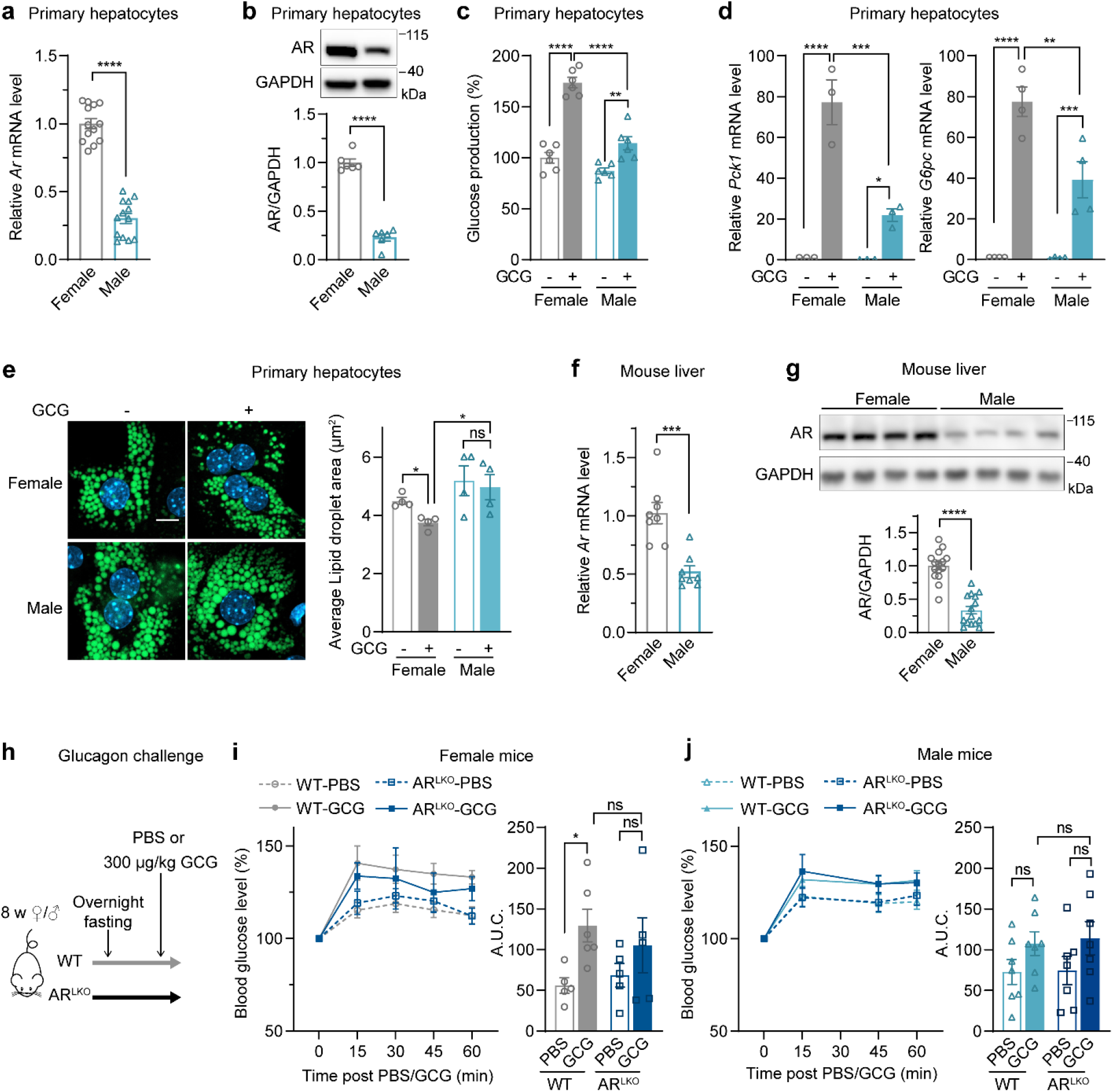
Higher expression of hepatic AR in female mice renders them more sensitive to glucagon treatment. **a, b,** The expression of AR in primary hepatocytes isolated from female and male littermates (n≥6 mice/group) was examined by qRT-PCR (**a**) and WB (**b**). The representative WB image (top) and quantification results (bottom) were shown. **c, d,** Hepatic glucose production (**c**) and qRT-PCR analysis of gluconeogenic genes *Pck1* and *G6pc* (**d**) in primary hepatocytes isolated from female and male littermates. The hepatocytes were treated with vehicle or 20 nM glucagon for 18 hours. **e,** The lipid droplets in hepatocytes were detected using BODIPY staining, and their sizes were quantified in primary hepatocytes obtained from female and male littermates. These hepatocytes were treated with either vehicle or 100 nM glucagon for a duration of 15 hours. Scale bars, 10 μm. (n>3000 lipid droplets/group using 80-120 hepatocytes from 4 mice). **f, g,** The expression of AR in the livers of female and male littermate mice (n≥8 mice/group) was examined using qRT-PCR (**f**) and WB analysis (**g**). The representative WB image (top) and quantification results (bottom) were shown. **h-j,** Glucagon challenge assay was conducted in both male and female WT and AR^LKO^mice (**h**). Blood glucose levels were measured before and after intraperitoneal administration of PBS or 300 μg/kg glucagon at specified time points in female WT or AR^LKO^ mice (**i**) and male WT or AR^LKO^ mice (**j**) (n≥5/group). The blood glucose of each mouse were normalized with basal blood glucose as 100%. The corresponding glucose curves were used to calculate the area under curve (A.U.C.), which is indicated adjacent to each curve. The data were presented as mean ± SEM from three or more independent experiments or from mice with the indicated number. The statistical significance was determined using the following criteria: ****p≤0.0001, ***p≤0.001, **p≤0.01, *p≤0.05; ns denotes no significance with p > 0.05. The statistical tests employed were Student’s t-test for the bar graph (**a, b, f, g**) and two-way ANOVA for (**c-e, i, j**).

To investigate the impact of hepatic AR on glucagon sensitivity *in vivo*, we generated liver-specific AR knockout (AR^LKO^) mice (Extended Data Fig. 4a). Hepatic AR deficiency had no effect on the body weights of male and female mice (Extended Data Fig. 4b, c). To evaluate the glucagon response, we conducted a glucagon challenge in eight-week-old littermates with either a wild type (WT) or AR^LKO^ genetic background, comprising both sexes (Fig. 2h). The results demonstrated a significant increase in blood glucose levels in female WT mice upon glucagon treatment (Fig. 2i). Surprisingly, the glucagon-induced rise in blood glucose levels was compromised in female AR^LKO^ mice. In contrast, no significant differences in response to glucagon administration compared to PBS injection were observed between male WT and AR^LKO^ mice (Fig. 2j). These findings strongly indicate that hepatic AR expression levels play a vital role in mediating the impact of glucagon on gluconeogenesis outcomes in female mice.

### Estrogen-related receptor alpha (ERRα) acts downstream of the non-canonical AR signaling in response to glucagon

AR plays a critical role as a ligand-activated transcription factor in the regulation of target gene expression. In addition to its canonical function, AR also exerts non-canonical functions by interacting with other transcription factors or forming complexes in prostate cancer cells^39,40^. We thus investigated the potential of the synthetic androgen methyltrienolone (R1881) to activate AR and enhance glucagon-stimulated glucose production by promoting the translocation of AR from the cytosol to the nucleus. The results showed that R1881 successfully induced the nuclear localization of cytosolic AR (Extended Data Fig. 5a). However, it did not result in elevated hepatic glucose production or the upregulation of genes associated with gluconeogenesis (Extended Data Fig. 5b, c). Notably, R1881 significantly attenuated the stimulatory effect of glucagon on glucose production (Extended Data Fig. 5b). These findings suggest that the non-ligand-binding or non-canonical AR may play a crucial role in mediating the activity of glucagon.

To unbiasedly explore the downstream elements responsible for AR-mediated glucagon activity, we conducted an RNA sequencing (RNA-seq) assay on female mouse hepatocytes treated with *Ar* knockdown and glucagon, along with appropriate controls (Fig. 3a). Among the 14,863 hits detected, there were 2,123 differentially expressed genes (fold change >2, false discovery rate <0.05) in primary hepatocytes, including 1,135 downregulated and 988 upregulated by the *Ar* knockdown (Fig. 3a). Furthermore, a group of genes consisting of 840 downregulated genes and 794 upregulated genes were identified in the *Ar* knockdown group, regardless of glucagon treatment (Fig. 3a and Supplementary Table 2). Subsequent analysis of transcription factor enrichment, using the 840 genes that exhibited a positive correlation with the AR expression at both basal and glucagon stimulated conditions, highlighted the nuclear hormone receptor ERRα as the most significant hit (Fig. 3b). To the best of our knowledge, there have been no previous studies investigating the role of ERRα in AR activity and the glucagon response.

**Fig. 3.**
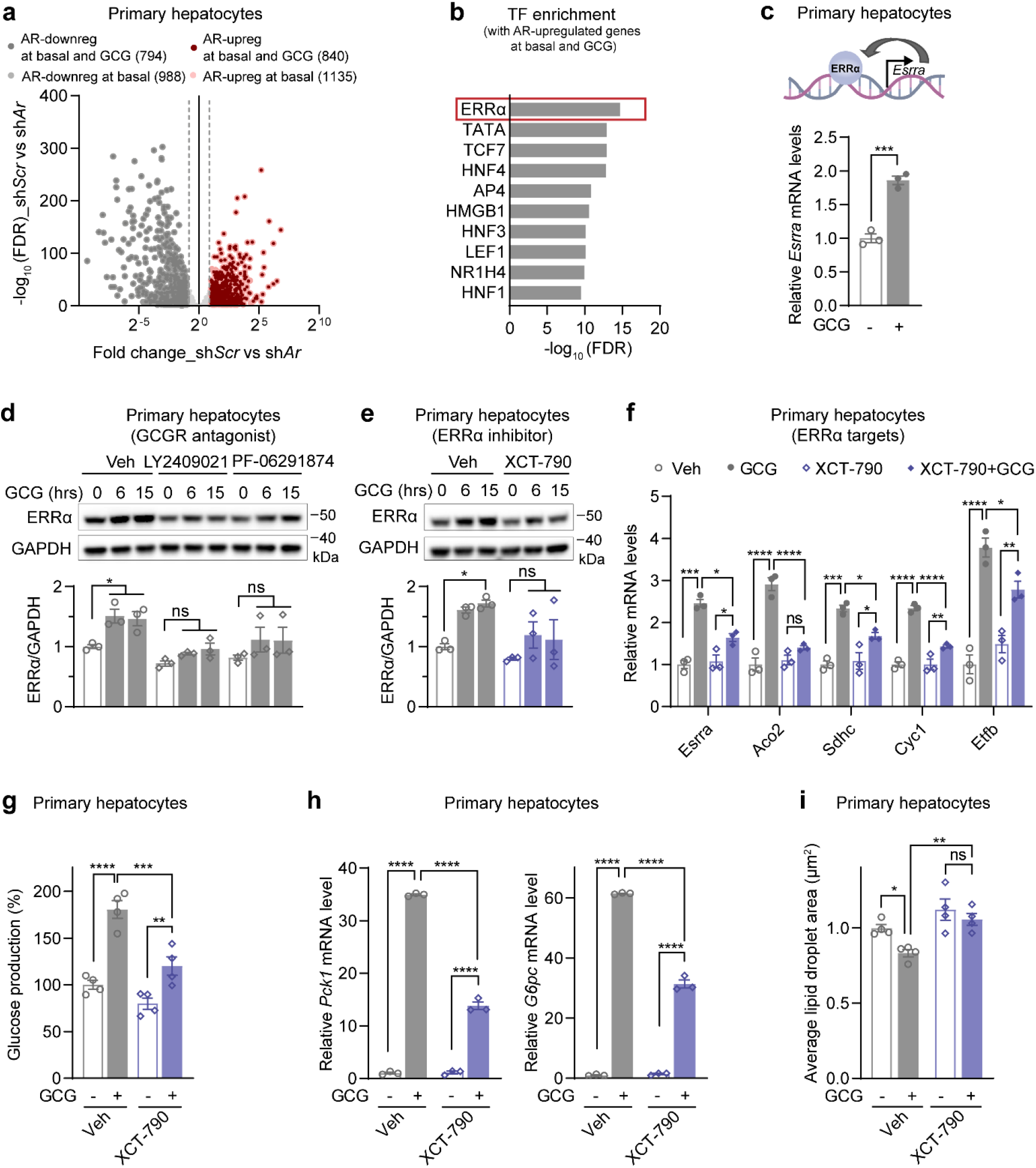
ERRα plays a crucial role in the regulation of hepatic gluconeogenesis and lipid breakdown during glucagon treatment. **a,** The volcano plot for the comparison of gene expression between Ad-sh*Scr* and Ad-sh*Ar* infected female mouse hepatocytes. Transcripts that showed significant differential expression (fold change (FC) > 2, false discovery rate (FDR) < 0.05) and were regulated by AR were colored in pink or grey. Additionally, transcripts that were similarly regulated by AR under glucagon-stimulated conditions are further highlighted in dark red or dark grey. **b,** A total of 840 AR-upregulated genes, which were identified under both basal and glucagon-stimulated conditions (colored in dark red), were subjected to further analysis using transcription factor enrichment in the MSigDB database. **c,** The expression of *Esrra* in female mouse primary hepatocytes treated with vehicle or 100 nM glucagon for 15 hours was analyzed using qRT-PCR (bottom). The above illustration displayed an established auto-regulatory loop regulation of ERRα activity. **d, e,** The ERRα protein levels in primary hepatocytes isolated from female mice were examined by WB analysis. The hepatocytes were pretreated with Veh or indicated compounds (2.5 μM LY2409022 (**d**), 20 μM PF-06291874 (**d**), and 2 μM XCT-790 (**e**) respectively) for 4 hours before stimulation with 100 nM glucagon for indicated time. The representative WB image (top) and the corresponding quantification results (bottom) were presented. **f,** The expressions of ERRα target genes *Esrra*, *Aco2*, *Sdhc*, *Etfb*, and *Cyc1* were examined by qRT-PCR in female mouse primary hepatocytes pretreated with Veh or 2 μM XCT-790 4 hours before stimulation with vehicle or 100 nM glucagon for 15 hours. **g, h,** Hepatic glucose production (**g**) and qRT-PCR analysis of the expression of gluconeogenic genes *Pck1* and *G6pc* (**h**) in female mouse primary hepatocytes pretreated with Veh or 2 μM XCT-790 4 hours before treatment with vehicle or 20 nM glucagon for 18 hours. **i,** The presence of the hepatocyte lipid droplets was detected using BODIPY staining, and their sizes were quantified in female mouse primary hepatocytes. These hepatocytes were pretreated with either Veh or 2 μM XCT-790 4 hours prior to treatment with vehicle or 100 nM glucagon for 15 hours (n>3000 lipid droplets/group using 80-120 hepatocytes from 4 mice). The data were presented as mean ± SEM from three or more independent experiments. The statistical significance was determined using the following criteria: ****p≤0.0001, ***p≤0.001, **p≤0.01, *p≤0.05; ns denotes no significance with p > 0.05. The statistical tests employed were Student’s t-test for the bar graph (**c**) and two-way ANOVA for (**d-i**).

The transcription factor ERRα is known to employ a feedforward regulatory mechanism that upregulates both transcription and protein expression of its own gene^41,42^. To assess whether glucagon administration could activate ERRα and determine if this activation could be mitigated by glucagon receptor antagonists LY2409021 and PF-06291874, we employed a readout involving this feedforward outcome. The results showed that glucagon treatment significantly upregulated the expression of ERRα, at both the mRNA and protein levels (Fig. 3c, d). Furthermore, the administration of either the glucagon receptor antagonists LY2409021 and PF-06291874, or the ERRα inhibitor XCT-790 markedly mitigated the effect of glucagon on ERRα expression (Fig. 3d, e). Notably, glucagon administration resulted in a significant upregulation of canonical ERRα target genes, such as *Esrra* itself, the aconitase 2 (*Aco2*), succinate dehydrogenase complex, subunit C (*Sdhc*), cytochrome c-1 (*Cyc1*) and the electron transfer flavoprotein subunit beta (*Etfb)* (Fig. 3f). Moreover, co-administering XCT-790 markedly attenuated all observed effects in primary hepatocytes following glucagon treatment, including the induction of ERRα target gene expression (Fig. 3f), glucose production (Fig. 3g), gluconeogenesis gene expression (Fig. 3h), and lipid droplet size outcomes (Fig. 3i). Together, these findings demonstrate the necessity of ERRα activation for the glucagon-mediated regulation of glucose and lipid metabolism in hepatocytes.

### Hepatic AR is required for the activation of ERRα by glucagon

Using changes in ERRα protein levels and mRNA expression of its target genes as experimental readouts, we assessed the correlation between hepatic AR and glucagon-stimulated ERRα activation. The results revealed that inhibition of AR, either chemically (Fig. 4a, b) or genetically (Fig. 4c, d), attenuated the upregulation of both ERRα protein levels and its downstream target gene expression induced by glucagon (Fig. 4a-d). Notably, we observed higher expression of AR in female mouse hepatocytes and liver compared to males (Fig. 2 a, b, f, g), potentially conferring increased sensitivity to glucagon (Fig. 2c-e, i, j). Moreover, we investigated whether glucagon similarly induces the expressions of ERRα and its associated target genes, implying heightened glucagon sensitivity. Strikingly, the results consistently showed significant and robust increases in the expressions of ERRα and its associated target genes in female mouse hepatocytes (Fig. 4e, f). To further investigate the role of ERRα in mediating the effecct of AR on glucagon functions, we utilized adenoviral techniques to overexpress ERRα in primary hepatocytes (Extended Data Fig. 6). Subsequently, we evaluated whether these cells could alleviate enzalutamide’s inhibitory effects on glucagon-mediated glucose production. Our findings demonstrate that the overexpression of ERRα effectively counteracts enzalutamide’s suppression of glucagon-induced glucose production (Fig. 4g). Collectively, these findings suggest that the activation of ERRα is mediated by AR following glucagon treatment. Subsequently, the activated ERRα modulates gluconeogenesis and lipid catabolism in response to glucagon.

**Fig. 4.**
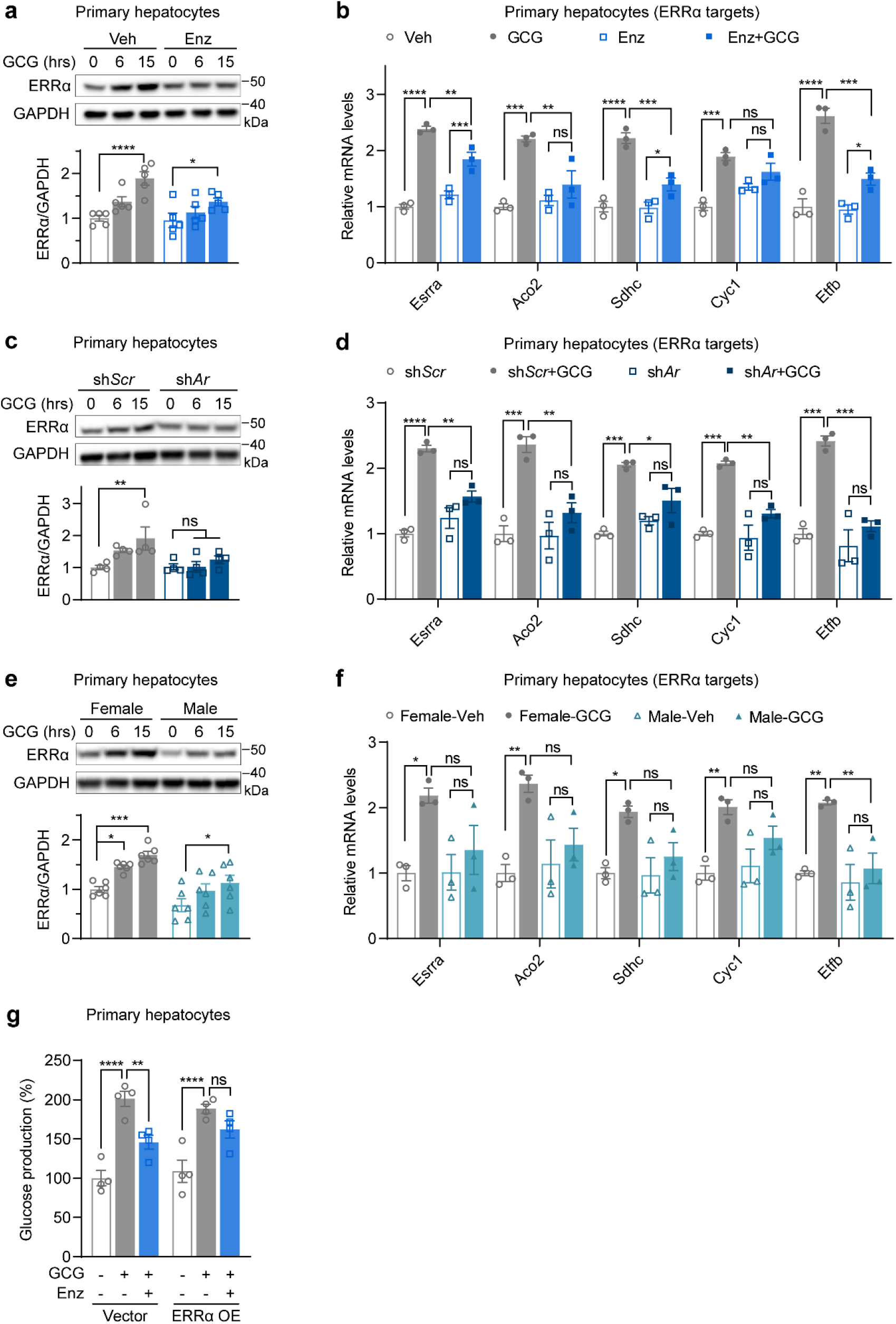
Hepatic AR acts through ERRα to modulate the activity of glucagon. **a, b,** The ERRα protein levels (**a**) and the expression of its representative target genes (**b**) were measured in female mouse primary hepatocytes pretreated with Veh or 20 μM Enz for 4 hours, followed by 100 nM glucagon stimulation for indicated time (**a**) or 15 hours (**b**). The representative WB image (top) and quantification results of ERRα protein (bottom) were shown. **c, d,** The ERRα protein levels (**c**) and the expression of its representative target genes (**d**) were measured in female mouse primary hepatocytes infected with Ad-sh*Scr* or Ad-sh*Ar* for 30 hours, followed by 100 nM glucagon stimulation for indicated time (**c**) or 15 hours (**d**). The representative WB image (top) and quantification results of ERRα protein (bottom) were shown. **e, f,** The ERRα protein levels (**e**) and the expression of its representative target genes (**f**) were measured with primary hepatocytes of female and male littermates. The hepatocytes were treated with vehicle or 100 nM glucagon for the indicated time (**e**) or 15 hours (**f**). The representative WB image (top) and quantification results of ERRα protein (bottom) were shown. **g,** The female mouse hepatocytes overexpressing an empty vector or ERRα were pretreated with either Veh or 20 μM Enz for 4 hours, followed by treatment with vehicle or 20 nM glucagon for 18 hours, hepatic glucose production was measured. The data were presented as mean ± SEM from three or more independent experiments. The statistical significance was determined using the following criteria: ****p≤0.0001, ***p≤0.001, **p≤0.01, *p≤0.05; ns denotes no significance with p > 0.05. The statistical tests employed were two-way ANOVA for (**a-g**).

### AR promotes the activation of ERRα by facilitating the interaction between PGC1α and ERRα

Peroxisome proliferator-activated receptor gamma coactivator 1-alpha (PGC1α) is a downstream effector of glucagon signaling^29,43^ and a potent coactivator of ERRα^20,44^. Previous studies have indicated that AR interacts with PGC1α in specific cancer cell lines^45^. Therefore, we hypothesized that AR could activate ERRα transcriptional activity by facilitating the interaction between ERRα and PGC1α (Fig. 5a). To test this hypothesis, we conducted immunoprecipitation (IP) experiments to examine the interactions between PGC1α, ERRα, and AR. Specifically, we overexpressed PGC1α and assessed its interactions with AR and ERRα in hepatocytes from WT or AR^LKO^ mice, under both glucagon stimulation and non-stimulated conditions. Our results revealed a significant increase in the interaction between PGC1α and ERRα in response to glucagon treatment in wild-type cells. However, this effect was not observed in AR knockout cells (Fig. 5b). Furthermore, we sought to investigate the influence of AR expression levels on the interaction between PGC1α and ERRα. To accomplish this, we performed IP experiments and ERRα activity tests in HEK 293T cells. The co-expression of AR resulted in an increased interaction between PGC1α and ERRα, with this interaction being directly proportional to the level of AR expression, suggesting a dose-dependent relationship (Fig. 5c). Notably, the administration of enzalutamide compromised AR’s ability to promote ERRα/PGC1α interaction (Fig. 5c). Additionally, we assessed the ERRα transcriptional activity in HEK 293T cells by overexpressing ERRα and PGC1α and utilizing the pyruvate dehydrogenase kinase 4 (*Pdk4*) luciferase reporter. Notably, the co-expression of ERRα and PGC1α, rather than the individual expression of either factor, significantly increased the transcriptional activity of the *Pdk4* promoter (Fig. 5d). These findings align with a previous study conducted by Wende *et al*^19^. Moreover, overexpression of AR further augmented the transcriptional activity of ERRα; however, this effect was hindered when enzalutamide was administered simultaneously (Fig. 5d). Collectively, these results suggest that AR activates ERRα by promoting its interaction with PGC1α.

**Fig. 5.**
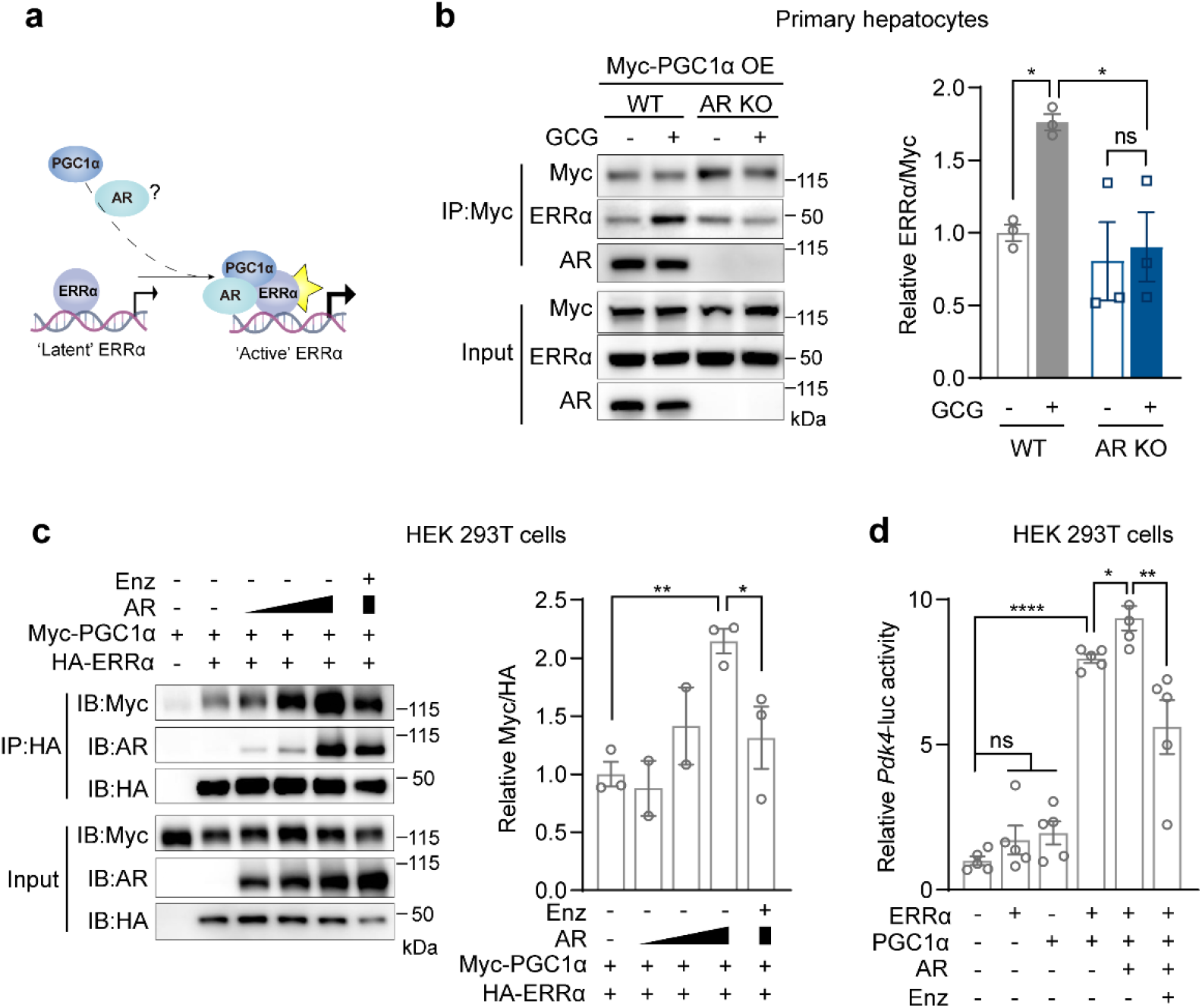
AR induces the activation of ERRα by facilitating its interaction with PGC1α. **a,** A hypothetical illustration for AR-mediated ERRα activation involving PGC1α. **b**, Western blot analysis was performed on the IP samples obtained from primary hepatocytes isolated from female WT and AR^LKO^ mice. The left side of the figure displays a representative western blot image, while the right side presents the corresponding quantification results. **c,** The interaction between AR protein and PGC1α/ERRα was examined via immunoblot analysis of the IP outcomes of HA-mERRα in HEK 293T cells. These cells were overexpressed with HA-tagged mERRα and Myc-tagged mPGC1α, either without AR or with varying doses of AR, as well as with 20 μM Enz, the representative image (left) and the qantification result (right) were shown. **d,** The luciferase activity of the *Pdk4*-luc reporter in HEK 293T cells was assessed in the context of overexpression of ERRα, PGC1α, or the ERRα/PGC1α complex. The experiments were conducted under varying conditions, including the absence or presence of AR, and with or without the administration of 20 μM Enz. The data were presented as mean ± SEM from three or more independent experiments. The statistical significance was determined using the following criteria: ****p≤0.0001, **p≤0.01, *p≤0.05; ns denotes no significance with p > 0.05. The statistical test employed was Student’s t-test for (**c-d**) and two-way ANOVA for (**b**).

### Hepatic AR couples ERRα to regulate glucagon-stimulated mitochondrial respiration and lipid breakdown

To elucidate the downstream effectors of the AR/PGC1α/ERRα signaling axis in response to glucagon treatment, we analyzed the ERRα target genes (GSE43638) annotated in a previous study^46^ that were also regulated by both AR and glucagon treatment. Remarkably, we observed a significant decrease in the expression of a cluster of genes encoding mitochondrial proteins, which are primarily involved in lipid metabolism and energy regulation through the tricarboxylic acid (TCA) cycle and oxidative phosphorylation (OXPHOS). This reduction occurred upon the depletion of AR by shRNA knockdown in primary hepatocytes (Fig. 6a). Notably, approximately half of these AR-downregulated genes demonstrated increased expression after glucagon treatment, suggesting that glucagon might regulate mitochondrial function by inducing the expression of ERRα targets.

**Fig. 6.**
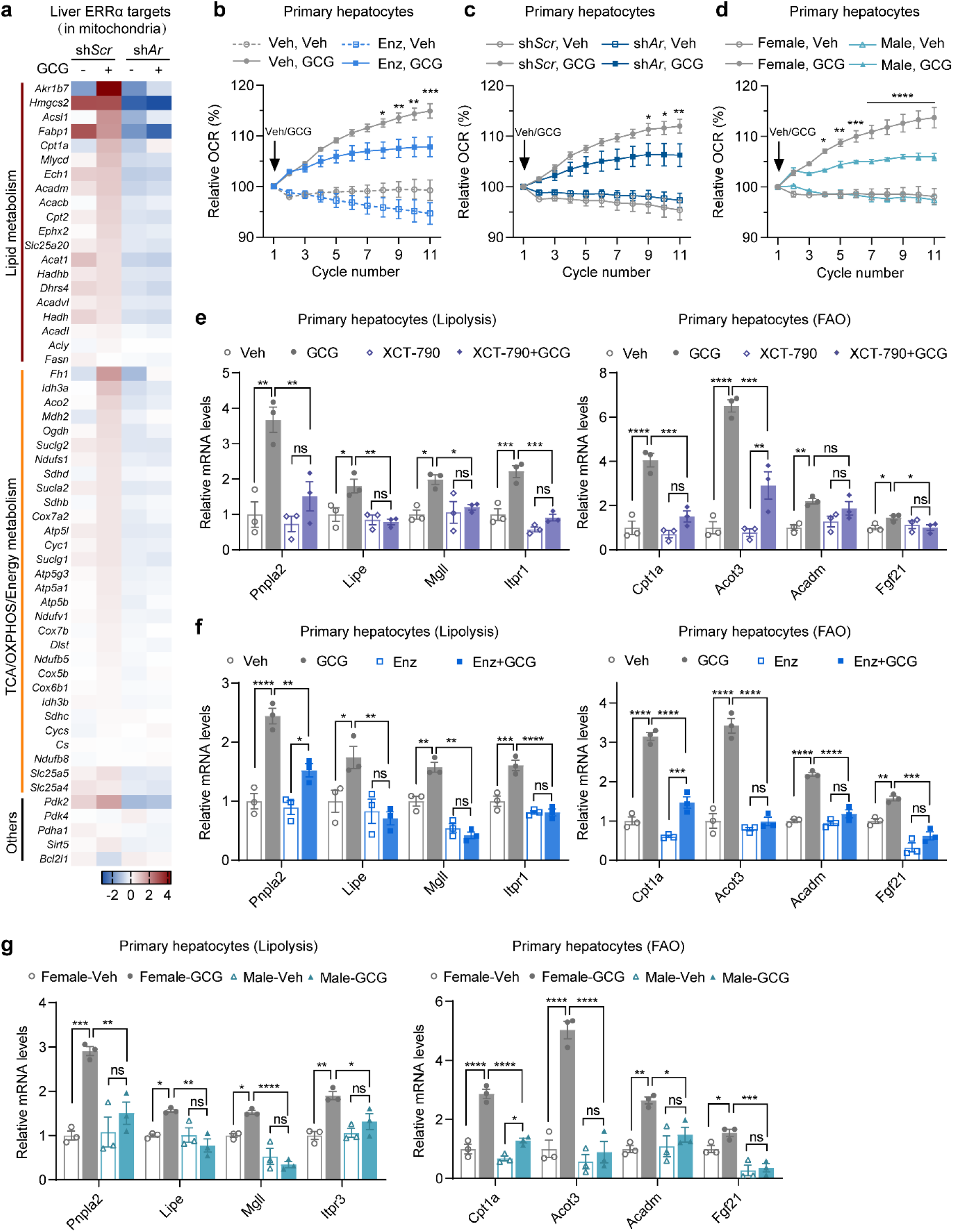
The hepatic AR and ERRα signaling cascade functions by regulating mitochondrial respiration and lipid catabolism in response to glucagon. **a,** A heatmap depicting ERRα target genes localized in mitochondria and clustered based on their metabolic functions. For this analysis, RNAseq was conducted on female mouse primary hepatocytes infected with Ad-sh*Scr* and Ad-sh*Ar*, and subsequently treated with either Veh or glucagon. **b, c,** The basal oxygen consumption rate (OCR) was measured following the injection of either Veh or 200 nM glucagon during the Seahorse assay. These female mouse primary hepatocytes were pretreated with either Veh or 30 μM Enz for a duration of 20 hours (**b**). Additionally, the cells were infected with either Ad-sh*Scr* or Ad-sh*Ar* (**c**). The OCR values were normalized to the corresponding first cycle within each group, with a value of 100%. d, The OCR test for primary hepatocytes of male and female littermates. These hepatocytes were treated with either Veh or 200 nM glucagon during the Seahorse assay. The OCR values were normalized to the corresponding first cycle within each group, with a value of 100%. **e, f,** The expression of lipolysis and fatty acid oxidation (FAO) genes were assessed using qRT-PCR in primary hepatocytes isolated from female mice. The hepatocytes were pretreated with either Veh or 2 μM of XCT-790 (**e**) or 30 μM of Enz (**f**) 4 hours prior to stimulation with either the vehicle or 100 nM of glucagon for 15 hours. **g,** The expression of lipolysis and fatty acid oxidation (FAO) genes were assessed using qRT-PCR in primary hepatocytes of female and male littermates. The hepatocytes were treated with vehicle or 100 nM glucagon for 15 hours. The data were presented as mean ± SEM from three or more independent experiments. The statistical significance was determined using the following criteria: ****p≤0.0001, ***p≤0.001, **p≤0.01, *p≤0.05; ns denotes no significance with p > 0.05. The statistical tests employed were two-way ANOVA for (**b-g**).

To investigate the effects of glucagon on mitochondrial respiration, we conducted a Seahorse assay in primary hepatocytes. The results demonstrated a gradual and significant enhancement of basal mitochondrial respiration by glucagon, with no impact observed on maximal respiration capacity (Extended Data Fig. 7). These findings were consistent with the results of a recent study^4^. Next, we examined the impact of AR on glucagon-stimulated basal mitochondrial respiration. To evaluate its influence, both a chemical inhibitor and genetic knockdown of AR were employed. Both approaches revealed a decline in glucagon-stimulated basal mitochondrial respiration compared to controls (Fig. 6b, c). Moreover, a higher sensitivity to glucagon stimulation in terms of basal mitochondrial respiration was observed in hepatocytes obtained from female littermates as compared to male littermates (Fig. 6d). These results further support the critical role of AR as a regulator of glucagon sensitivity.

Consistent with our RNAseq data (Fig. 6a), we observed a significant increase in the expression of genes related to hepatic lipolysis and fatty acid oxidation following glucagon treatment (Fig. 6e). Intriguingly, this effect was almost completely abolished when cells were pretreated with the ERRα inhibitor XCT-790 (Fig. 6e) or the AR inhibitor enzalutamide (Fig. 6f). Furthermore, primary hepatocytes obtained from female littermates exhibited greater glucagon-induced hepatic lipolysis and fatty acid oxidation compared to males (Fig. 6g). In summary, our study revealed the regulatory role of hepatic AR in glucagon signaling, which governs both gluconeogenesis and lipid catabolism through the ERRα/PGC1α pathways (Fig. 7).

**Fig. 7.**
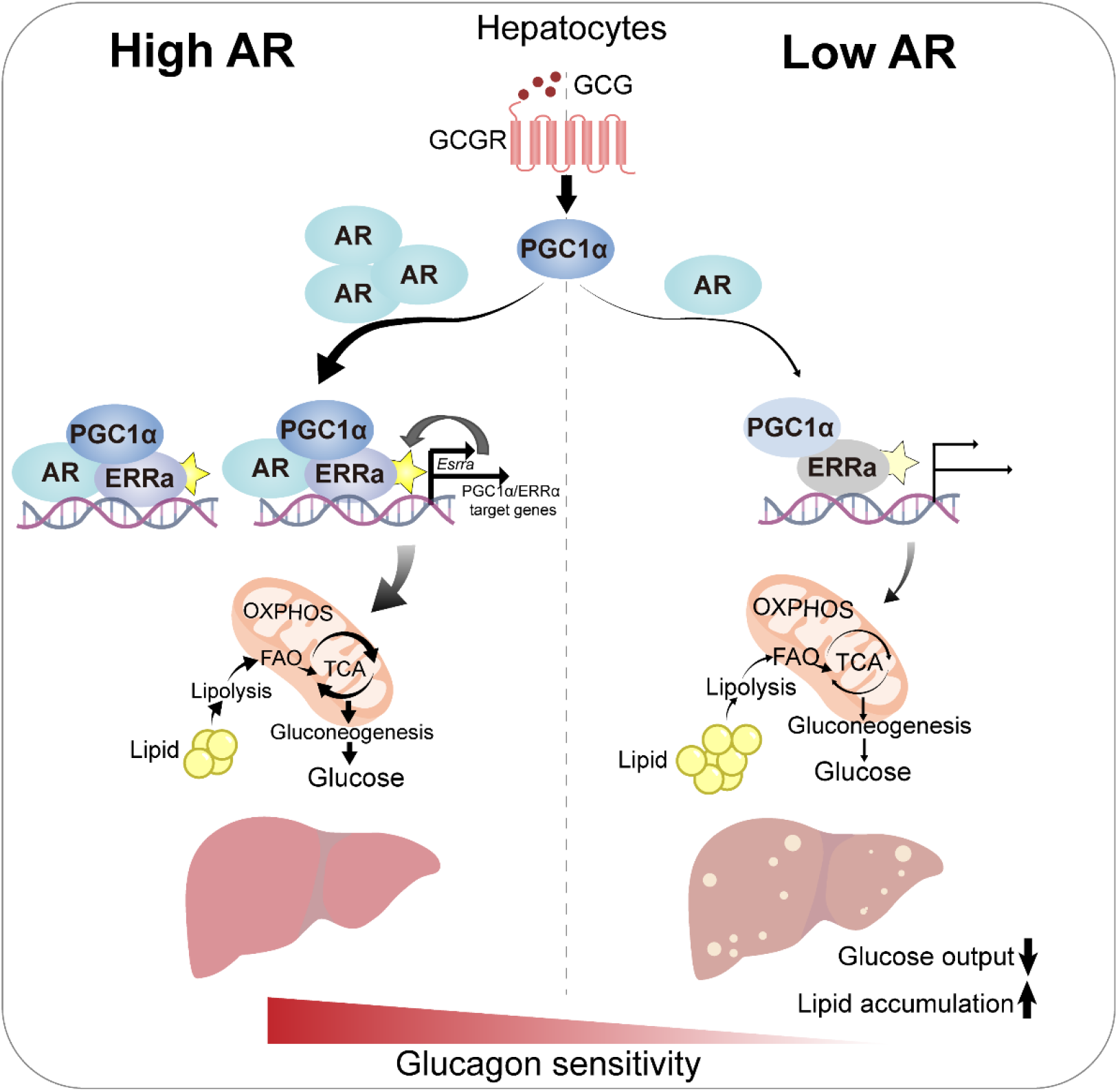
Working model for the action of hepatic AR in response to glucagon.

## Discussion

This study demonstrated that hepatic AR acts as a novel regulator of glucagon signaling. Specifically, we found a positive association between AR expression and glucagon sensitivity, as evidenced by enhanced activation of ERRα, hepatic lipolysis and fatty acid oxidation, mitochondrial respiration, and glucose production upon glucagon stimulation. Conversely, when AR was inhibited, glucagon signaling was impaired, leading to reduced glucose output and increased lipid accumulation in conditions of normal feeding (Fig. 7). Previous research has also highlighted the diverse metabolic functions performed by hepatic AR in various pathological states, including chronic androgen treatment^14,47^ or high-fat diet feeding conditions^16^. Mechanistic analysis of the current study revealed that the AR functions via the PGC1α/ERRα/mitochondria axis to regulate glucagon-stimulated hepatic lipid catabolism and gluconeogenesis in normal physiology, primarily in female mice. Overall, these findings contribute significantly to our understanding of the role played by hepatic AR in coordinating glucose and lipid metabolism in response to glucagon.

Our chemical screen identified that enzalutamide, an AR inhibitor, can upregulate the expression of CeACAD10 in *C. elegans*. Despite the absence of a clear AR homolog with high sequence similarity to humans in *C. elegans*, the organism does possess various nuclear receptors that have not been extensively characterized^48^. Some evidence suggests that functional AR homologs may exist in *C. elegans*. Furthermore, a previous study has reported that androgen treatment can impact the behavior of *C. elegans*^49^. Our study revealed that, apart from enzalutamide, other androgen receptor inhibitors in our drug library also induced the activity of CeACAD10 at a concentration of 100 μM (data not shown). However, further investigation is required to ascertain which nuclear receptor fulfills the role of AR in *C. elegans*.

Our findings indicate that both enzalutamide and metformin have the capacity to decrease glucagon-stimulated gluconeogenesis in primary hepatocytes. However, unlike metformin, enzalutamide does not demonstrate comparable effects in reducing blood glucose levels *in vivo*. This disparity can be attributed to the distinct mechanisms of action employed by these two drugs, although they ultimately intersect at mitochondrial function. Specifically, metformin is widely recognized as a direct inhibitor of mitochondrial complex I^50–52^, whereas our study reveals that enzalutamide acts through the AR/PGC1α/ERRα signaling axis to modify mitochondrial function and lipid catabolism.

Canonically, AR functions by translocating its androgen-bound form to the nucleus through a genomic or transcriptional pathway. Once in the nucleus, AR acts as a direct regulator of gene transcription by binding to specific target promoters^53^. Our results showed that the inhibition of AR activity by enzalutamide or flutamide can decrease glucagon-stimulated gluconeogenesis. It has been well-documented that these compounds can directly bind to AR and hinder its nuclear translocation^34,54^. Additionally, our findings indicate that androgen stimulation significantly enhances nuclear localization of AR in primary hepatocytes. Surprisingly, however, this stimulation does not enhance glucagon-stimulated glucose production; instead, it even reduces it. Thus, the mechanisms by which AR mediates glucagon signaling may not involve its typical cytosol-nucleus translocation activity. Instead, our results suggest that the availability of free AR in the cytosol plays a crucial role in glucagon signaling transduction and its associated effects.

Our study has revealed that ERRα serves as a major surrogate for AR in the regulation of hepatic gene expression. This finding is supported by our transcription factor analysis, which demonstrated an enrichment of ERRα in AR-regulated genes. Mechanistically, we observed an interaction between AR and both PGC1α and ERRα, leading to an increase in the transcriptional activity of ERRα. This suggests that AR regulates glucagon sensitivity through a non-canonical way. However, further investigation is required to identify the precise location where AR promotes the binding between PGC1α and ERRα.

We observed that female mouse hepatocytes exhibited higher levels of gluconeogenesis in response to glucagon compared to their male counterparts. This difference in glucagon sensitivity between sexes is likely attributed to the differential expression of hepatic AR. In fact, all downstream events related to the support of gluconeogenesis were more significant in female mouse hepatocytes than in their male littermates. These events included glucagon-stimulated ERRα transcriptional activity, mitochondrial respiration, and lipid catabolism. Furthermore, it has been noted that the prevalence of NAFLD is higher in men than in women^55,56^. It is also worth noting that impaired glucagon signaling has been associated with an increased risk of NAFLD in patients with type 2 diabetes^2,3^. Additionally, patients with spinal-bulbar muscular atrophy who have AR mutations have a higher prevalence of NAFLD^57,58^. However, the underlying mechanisms behind this increased NAFLD prevalence remains unclear. Our study provides a potential mechanistic link that may explain these clinical observations, which could involve impaired glucagon signaling due to the loss or reduction of AR function in the liver. Further research is needed to establish this link experimentally. It would be also worthwhile to investigate whether therapies that enhance AR signaling, either by increasing AR expression or activating ERRα, can improve glucagon sensitivity and halt the progression of NAFLD.

One notable limitation of this study is the absence of validation regarding the roles of PGC1α and ERRα in glucagon activity in mice, despite their demonstrated effects in isolated primary hepatocytes. Furthermore, the impact of androgen levels on the observed sex differences in glucagon sensitivity has not been explored beyond the expression level of AR. Additionally, our analysis of the disparities in glucagon response between sexes is confined to mouse littermates aged 6-16 weeks. To increase the credibility of our findings, future research should investigate variations in AR expression and glucagon response across different age groups.

## Acknowledgments

We thank Drs. Yan Wang, Xuedan Li, Shang Cai, Yuxiong Feng, Zhenji Gan, Yifu Qiu for kindly providing reagents for this study. We also thank Drs. Ben Zhou, Zhenji Gan, Yong Liu, Min Jiang, Chunqing Song, Haojie Yu and Wenjing Su for their invaluable support and discussions. This work was supported by the National Natural Science Foundation of China (32071151 and 32271357 to L.W; 32100949 to J.Y.), the Natural Science Foundation of Zhejiang Province (2022XHSJJ005 to L.W.), and the HRHI program (202209008 to L.W.) of Westlake Laboratory of Life Sciences and Biomedicine.

## Author Contributions

J.C. and J.Y. conducted the cell and mouse experiments and analyzed the data. Y.W. and J.C. conducted compound screen in *C. elegans*. W.H. established culture system for *C. elegans* and analyzed the expression of ERRα targets in primary hepatocytes treated with drugs. L.W., J. Y. and J.C. designed the experiments and wrote the manuscript with all inputs of others.

## Methods

### Compound screen in *C. elegans*

The *C. elegans* stain MGH249 *alxls19 [CeACAD10p::CeACAD10::mRFP3-HA, myo-20::GFP]*^32^ was maintained with *Escherichia coli* OP50 and maintained at 20 ℃ on nematode growth media (NGM) plates. For compound screen, synchronized worms at L1 stage were dropped onto 96-well NGM plates (∼50 worms per well) that were seeded with the *E. coli* OP50 bacteria and the chemicals 4 hours before dropping worms. All chemicals (TargetMol) used in the screen were dissolved in DMSO and applied at 50 μM. 56 hours after chemical treatment, the worms at young adult stage were washed off from cultured plates, subjected to 2 mg/mL levamisole for paralization, and transferred to 96-well slides for imaging with Leica DM6000 microscope. An arbitrary score was given to each image based on the fluorescence intensities of tested worms, with Veh-treated worms as score 0, the brightest as the score 5 and the darkest as the score −2. In the validation experiments, 3.5 cm NGM plates were utilized. The worm’s drug treatment strategies were comparable to those described above, with the exception that 50 mM metformin was incorporated for the purpose of comparison.

### Mice

The *Ar*-flox mice and Alb-Cre mice on C57BL/6 background were purchased from GemPharmatech (Nanjing, China). Liver-specific AR knockout mice were generated by crossing *Ar*-flox mice with Alb-Cre mice. C57BL/6J mice were used for primary hepatocytes isolation and the *in vivo* experiments. For experiments with sex difference comparison, the female and male littermates were from same breeders. For other experiments, C57BL/6J mice were purchased from Shanghai Jihui Laboratory Animal Care or Shanghai SLAC Laboratory Animal Center. All the mice were housed in SPF-level laboratory animal room of Westlake University on a 12/12 h light/dark cycle between 22-24 °C with ad libitum access to water and normal chow diet before experiments. Animal protocols were approved by the Westlake University Institutional Animal Care and Use Committee and conducted accordingly.

### Isolation and culture of primary hepatocytes

Primary hepatocytes were isolated from 7-12 week-old female and male mice with modified two-step collagenase perfusion method as described previously^59^. Briefly, liver was firstly perfused with 50 mL HBSS buffer supplemented with 0.5 mM EGTA, followed by perfusion with 50 mL HBSS containing 5 mM CaCl_2_ and 0.5 mg/mL collagenase type IV. The released liver cells were washed with cold M199 medium, filtered with 70 μm filter, and centrifuged at 50 g for 5 min at 4 °C. The cells were resuspended with 20 mL cold M199 medium. 20 mL percoll was added for gradient centrifugation at 270 g for 5 min at 4 °C. The hepatocytes precipitated at the bottom were washed once with cold M199 medium and resuspended with fresh M199 medium. Cells with the survival rate greater than 90% by trypan blue exclusion test were used in subsequent experiments. The primary hepatocytes were seeded on 5 μg/cm^2^ rat tail collagen type I coated 12- or 6- well plates at 2.5×10^5^ cells/mL in complete culture medium (M199 supplemented with 7.5% FBS, 1% penicillin/streptomycin, and 100 nM dexamethasone). After 4-5 hours attachment, the media was replaced with fresh medium for further experiments.

### Cell culture

HEK 293A cells and HEK 293T cells were cultured with high glucose DMEM supplemented with 10% FBS, 1% penicillin/streptomycin and incubated at 37 °C, 5% CO2 condition.

### Hepatic glucose production assay

The primary hepatocytes were cultured in serum-deprived medium overnight after allowing 4-5 hours for attachment in a complete culture medium following isolation. This was done prior to conducting glucose production assays on the second day. If the experiments involved adenovirus infection, the hepatocytes were replaced with fresh complete culture medium and cultured for an additional 24 hours. Subsequently, serum deprivation was performed overnight prior to glucose production assays on the third day.

The cells were washed once and placed in a glucose production medium for the glucose production assay. This medium was consisted of a phenol red-free DMEM with no glucose, supplemented with 2 mM pyruvate, 10 mM L-glutamine, 20 mM L-lactate, 25 mM Hepes, and 100 nM dexamethasone. Those cells were then stimulated with or without glucagon (20 nM/100 nM) for a period of 6 hours, as indicated.

To detect the glucose released from hepatocytes, the medium was collected and analyzed using the Amplex Red/Glucose Oxidase kit. Protein lysates were collected using 1% SDS and quantified using the BCA protein assay. Glucose production was normalized to total protein and expressed as untreated basal glucose production, with a value of 100%.

### Adenovirus plasmid construction and infection

The Ad-sh*Scr* and Ad-sh*Ar*#1/#2 were purchased from Hanbio Biotechnology, Shanghai. The target sequences were listed in Supplementary Table 3. The Ad-Vector, Ad-*Esrra*, and Ad-Myc-PGC1α plasmids were created in our laboratory. Mouse liver *Esrra* cDNA and Myc-PGC1α were cloned into pShuttle-CMV vector. The pShuttle-CMV, pShuttle-CMV-*Esrra* and pSuttle-CMV-Myc-PGC1α were linearized and transformed into BJ5183 competent cells containing Ad-Easy vector. Pac I-digested recombinant Ad plasmid DNA was transfected into HEK 293A cells using lipo8000 transfection reagent. After 7-10 days, when the cells exhibited significant cytopathic effect (CPE), they were collected and frozen in liquid nitrogen and thawed in a 37 °C water bath for more than 4 cycles to prepare the primary stock vial. The virus was then amplified through infection of HEK 293A cells for 2-3 generations, and its concentration was determined.

For the experiments, primary hepatocytes were allowed to attach for 4-5 hours in complete culture medium. Subsequently, the medium was replaced with fresh complete culture medium, and the cells were infected with adenovirus for a duration of 30 hours. Ad-sh*Scr* and Ad-sh*Ar*#1/#2 were used at a multiplicity of infection (MOI) of 5-10, while Ad-Vector and Ad-*Esrra* were used at an MOI of 30 and Ad-Myc-PGC1α at an MOI of 50.

### BODIPY staining

The isolated primary mouse hepatocytes were seeded on rat tail collagen type I coated glass coverslips in 24-well cell culture plates with complete culture medium. After 4-5 hours attachment, the cells were serum deprived and treated with vehicle or 30 μM enzalutamide overnight. Cells were stained with BODIPY 493/503 (8 μg/mL) for 15 min at 37 °C, followed by two times of wash with PBS and fixed with 4% paraformaldehyde for 30 min at room temperature. Cells were counterstained with 1 μg/mL DAPI for 5 min at room temperature, and mounted with mounting media for imaging. The images were acquired with Zeiss 800 equipped with Axiocam 702 mono camera. Images were analyzed with Zen 2.5 software. The size of lipid droplet (LD) was analyzed with Image J (Fiji). Briefly, the image type was firstly converted to 8-bit, followed by binary, watershed and LD size was measured with particle analysis set 1.5 µm^2^-infinity. For each group, more than 2000 lipid droplets were randomly picked and calculated the average lipid droplet area for further analysis.

### Oral gavage of enzalutamide in mice

Enzalutamide was prepared with vehicle solution of 5% DMSO, 1% (m/v) CMC (sodium salt of carboxymethyl cellulose) and 0.1% (v/v) Tween 80. 7-week-old female mice were randomly divided into two groups and administrated with vehicle or 30 mg/kg enzalutamide by oral gavage for 20-25 consecutive days. Body weights were recorded daily.

### Glucagon challenge assay

All mice were handled daily for several days before the commencement of the experiment to mitigate the potential impact of stress induced by handling during the course of the experiment. The mice were subsequently subjected to an overnight fasting period, followed by an intraperitoneal injection of either PBS or 300 μg/kg of glucagon. Blood glucose levels were measured using a glucometer (Accu-Chek) prior to the administration of the injection or at 15, 30, 45, and 60 min post-injection. For data analysis, the blood glucose of each mouse were normalized with basal blood glucose as 100%, and post glucagon injection glucose was analysed as the percentage change from baseline. For A.U.C. (area under curve) calculation, measure the area under the curve with the subtraction of the area under the baseline.

### Glucose tolerance test

Mice were fasted overnight and administrated with vehicle, 90 mg/kg enzalutamide or 150 mg/kg metformin by oral gavage, followed by the intraperitoneal injection of 1.5 g/kg glucose 1 hour later. Blood glucose levels were measured with glucometer (Accu-Chek) before or 15, 30, 60, and 120 min after injection of glucose.

### RNAseq analysis

The RNA-sequencing was performed by Novogene. Poly-A pull-down was used to enrich mRNA from total hepatocyte RNA. The purity of RNA was examined with the NanoPhotometer® spectrophotometer (IMPLEN) and quantity was determined by Qubit® RNA Assay Kit in Qubit®2.0 Flurometer (Life Technologies). RNA Nano 6000 Assay Kit of the Bioanalyzer 2100 system (Agilent Technologies) was used to evaluate the integrity of RNA. NEBNext® UltraTM RNA Library Prep Kit for Illumina® (NEB, USA) following manufacturer’s recommendations and index codes was used to generate sequencing libraries. The clustering of the index-coded samples was performed on a cBot Cluster Generation System using TruSeq PE Cluster Kit v3-cBot-HS (Illumia) according to the manufacturer’s instructions. After cluster generation, the library preparations were sequenced on an Illumina platform and 125 bp/150 bp paired-end reads were generated. Sequencing reads were demultiplexed (bcl2fastq) and quantified with kallisto 0.46 with default parameters to the Mouse GRCm38. Differential gene expression analysis was performed using DESeq2 1.26.0, and only false discovery rate (FDR)-adjusted P values less than 0.05 were considered statistically significant. The transcription factor enrichment was analyzed with genes of interest at MsigDB (http://www.gsea-msigdb.org/gsea/msigdb/annotate.jsp).

### Seahorse assay

The oxygen consumption was measured by Seahorse instrument as previously described^4^. Primary hepatocytes were seeded on collagen type I-coated XF96 seahorse cell culture plate (1×10^4^ cells/well) in complete medium for 4-5 hours, followed by replacing fresh medium and being infected with adenovirus (sh*Scr*, sh*Ar*) for 32 hours. The cells were then washed twice and incubated in serum-free medium with drugs overnight as indicated. The next morning, cells were washed with assay medium (DMEM base medium containing 1 mM pyruvate, 2 mM glutamine and 5.5 mM glucose, PH7.4). 200 μL assay medium containing different drugs was added to each well. Plates were equilibrated at 37 °C of non-CO_2_ incubator for 1 hour. Three measurements of basal oxygen consumption rates were recorded as baseline. After baseline measurements, vehicle or 200 nM glucagon was injected and the oxygen consumption was recorded. Ten measurements were taken following injection. Subsequently, all wells were injected with 2 μM oligomycin, 2 μM FCCP, and 1 μM antimycin A plus 1 μM rotenone to obtain different respiratory parameters. Oxygen consumption rates were calculated and presented with initial basal oxygen consumption rates as 100%.

### Luciferase assay

For the luciferase reporter assay, HEK 293T cells were co-transfected with the pRL-TK Renilla, *Pdk4* luciferase reporter and empty vector, Myc-PGC1α, ERRα or AR. Post 24 hours of transfection, cells were changed to fresh serum-deprived DMEM medium and treated with vehicle or enzalutamide for additional 20 hours. Cells were lysed and measured by using Dual Luciferase Reporter Gene Assay Kit (Beyotime) according to the kit instructions.

### Western blotting analysis

Cells were washed once with cold PBS and the total protein was extracted using RIPA lysis buffer supplemented with protease and phosphatase inhibitor cocktail. The whole cell lysates were centrifuged at 15000 g, 4 °C for 10 min and the supernatant was collected. For mouse tissue protein extraction, the tissues were homogenized in RIPA lysis with TissueLyser II (Qiagen), sonicated with Bioruptor sonication device (Diagenode) and spun down to collect supernatant. The protein was quantified by BCA protein assay. Loaded protein samples were separated with Bis-Tris polyacrylamide gels (GenScript) and transferred to nitrocellulose membrane. The membrane was blocked with 5% w/v non-fat milk, incubated with primary antibody overnight at 4 °C and secondary antibody at room temperature for 1 hour. The western blot signals were developed with chemiluminescent detection reagents and detected by GEL Imaging System (AI680). The signals were quantified with Image J.

### Immunoprecipitation

The primary hepatocytes were isolated from female WT and AR^LKO^ mice. These cells were infected with an adenovirus containing Myc-PGC1α (MOI=50) for 36 hours, followed by treatment with or without glucagon for 12 hours prior to the IP assay. In the IP assay conducted in HEK 293T cells, Myc-mPGC1α, HA-mERRα/HA-cDNA and Flag-mAR/Flag-mScarlet were transfected into cells for 24 hours. The cells were then treated with or without 20 μM enzalutamide for an additional 20 hours. Both hepatocytes and HEK 293T cells were lysed using NP-40 Lysis Buffer (Beyotime). After centrifugation at 15000 g, 4 °C for 10 min, about 5% of supernatant was aliquoted as the input, while the remaining was incubated with Myc-Nanab-Agarose or HA-Nanoab-Agarose beads for 4 hours at 4 °C to precipitate the target protein. The beads were washed three times with 1 mL of NP-40 Lysis Buffer and eluted with 1%SDS at 98 °C for 10 min. The eluate was then centrifugated at 2500 g, room temperature for 1 min, and the supernatant was collected for western blotting analysis.

### qRT-PCR

RNA was extracted using TRIzol according to manufacturer’s instructions. 500 ng RNA was reverse transcribed using HiScript III 1st Strand cDNA Synthesis Kit followed by qPCR reactions using SYBR Green master mix and analyzed by Biorad CFX Touch Real-Time PCR Detection System (SIA-PCR008). The primers used were listed in Supplementary Table 4.

### Immunofluorescence

The primary mouse hepatocytes were seeded on rat tail collagen type I coated glass coverslips in 24-well cell culture plates with complete culture medium. After 4-5 hours of attachment, the cells were serum deprived and treated with vehicle or 10 nM R1881 for 24 hours. The cells were washed twice with PBS, fixed with 4% paraformaldehyde for 15 min, permeabilized with PBS containing 0.5% Triton X-100 for 10 min at room temperature and blocked with 5% donkey serum for 1 hour in PBS buffer. Primary antibody against AR (1:100) was incubated overnight at 4 °C in the blocking solution. Next day, after washing with PBS three times, the slides were incubated with secondary antibody Alexa Fluor 488 donkey anti-rabbit IgG for 1 hour at RT, stained with 1 μg/mL DAPI for 3 min, rinsed with PBS 3 times, and mounted before imaging. Confocal images were acquired with Zeiss 800 equipped with Axiocam 702 mono camera. Images were analyzed with Zen 2.5 software.

### Liver triglyceride content

Triglycerides (TG) was extracted according to the protocol provided by the commercial kit. 20-30 mg liver tissue was homogenized in kit provided buffer by TissueLyser II (SIA-UH004). After centrifugation at 10000 g, 4 °C for 10 min, the supernatant was collected to measure TG concentration with Triglyceride Colorimetric Assay Kit. The pellets were boiled and fully dissolved in a solution containing 1% SDS before undergoing protein concentration determination. The TG content of each sample was normalized to its corresponding protein concentration.

### Oil Red O staining

The liver tissues were fixed in 4% paraformaldehyde for 2 hours, and dehydrated in 30% sucrose at 4 °C. Tissues were embedded in Tissue-Tek OCT compound and frozen at −80 °C. Then 10 μm liver sections were generated by cryostat and stained with Oil Red O stain kit according to the manufacturer’s protocol. Briefly, sections were washed with ddH_2_O and incubated with 60% isopropanol for 2 min, then stained with 0.5% Oil Red O for 10 min. Subsequently, sections were incubated for 2 min in 60% isopropanol and washed once with water. Finally, slides were counterstained with hematoxylin and mounted with Glycerol Jelly Mounting Medium. Olympus IX83 microscope was used to acquire images.

### Statistics

All statistical analyses were performed by using two-tailed unpaired Student’s *t* tests between two groups, one-way ANOVA between more than two groups or two-way ANOVA when there were more than two conditions using Prism 8.0. Data were presented as mean ± SEM, with the statistical significance differences indicated as: * *p* < 0.05; ** *p* < 0.01; *** *p* < 0.001; **** *p* < 0.0001 and ns denotes no significance with p > 0.05.

## Data Availability

The RNA-seq data have been made available at the Gene Expression Omnibus (GEO) repository under the accession number GSE236551.

## Figures and legends

**Extended Data Fig. 1.**
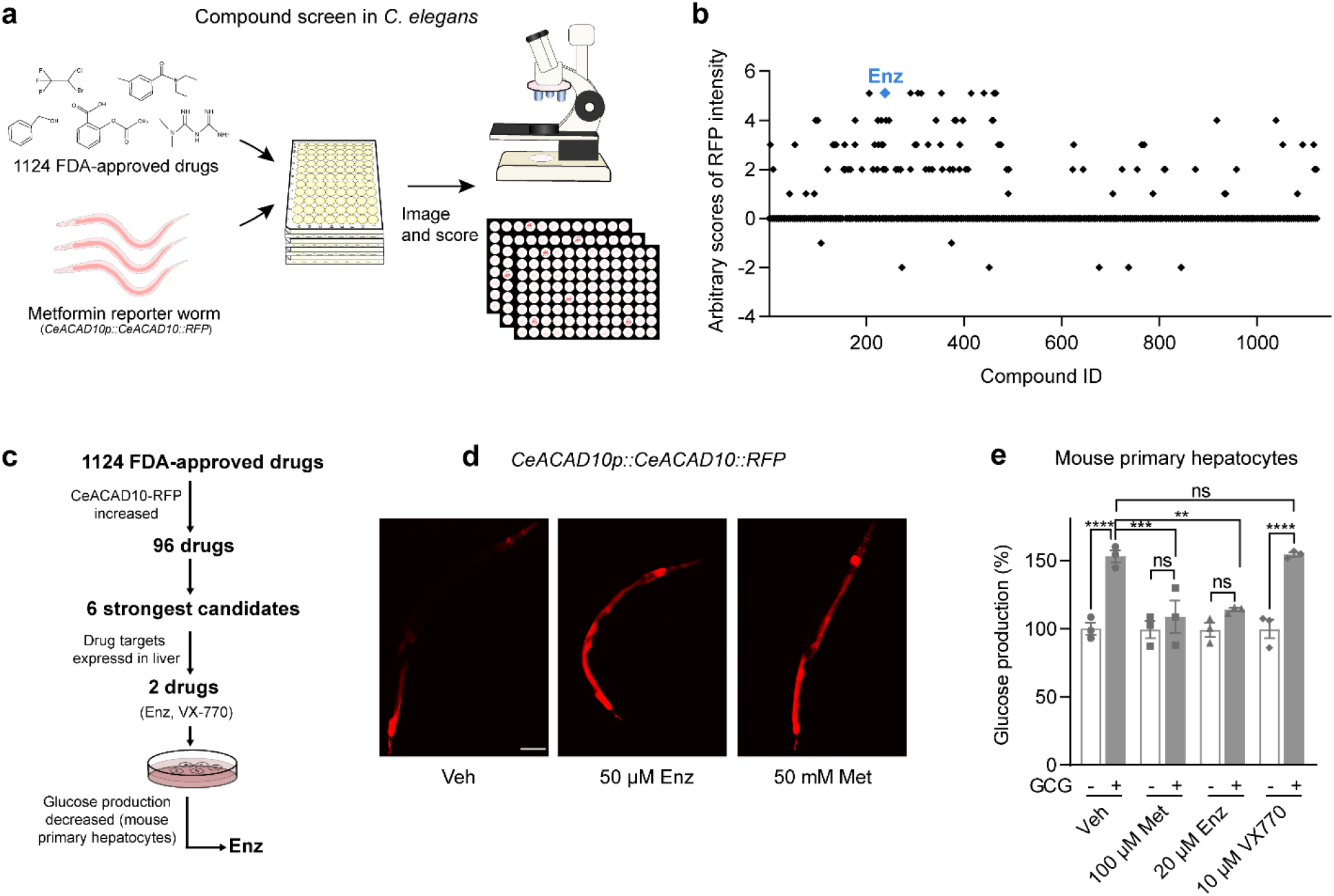
Compound screen for metformin mimickers in *C. elegans* and primary hepatocytes. **a,** The flow graph illustrates the compound screen strategy. **b,** Arbitrary scores for the 1,124 screened compounds, with the enzalutamide (Enz) compound highlighted. The X axis represents the compound ID in the library, while the Y axis represents the fluorescence intensity score of worms after treatment with each compound. **c,** A flowchart showing different filter conditions used in the screen. **d,** The representative images of CeACAD10-RFP worms treated with Veh, 50 μM Enz and 50 mM metformin (Met) for 56 hours. Scale bars, 100 μm. **e,** The glucose production of primary hepatocytes pretreated with Veh, 100 μM Met, 20 μM Enz, and 10 μM VX-770 respectively 4 hours prior to treatment with vehicle or 20 nM glucagon for another 18 hours. The data were presented as mean ± SEM from one or more independent experiments. The statistical significance was determined using the following criteria: ****p≤0.0001, ***p≤0.001, **p≤0.01; ns denotes no significance with p > 0.05. The statistical test employed was two-way ANOVA for (**e**).

**Extended Data Fig. 2.**
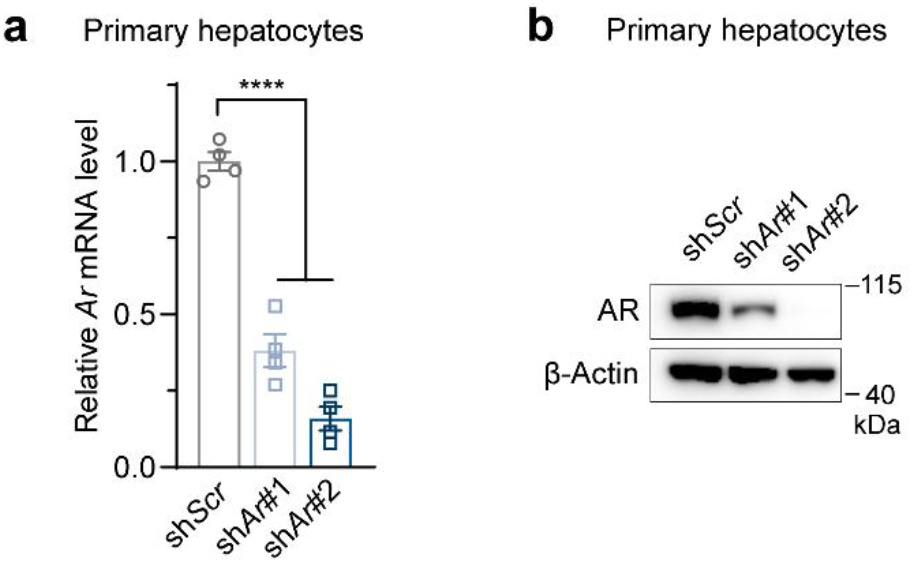
The knockdown efficiency of AR in primary hepatocytes treated with Ad-sh*Scr* or Ad-sh*Ar*. **a,b,** The expression of AR was examined in female primary hepatocytes post shRNA knockdown using qRT-PCR (**a**) and WB analysis (**b**). A representative WB image of three independent replicates was shown (**b**). The data were presented as mean ± SEM from three or more independent experiments. The statistical significance was determined using the following criteria: ****p≤0.0001. The statistical test employed was one-way ANOVA for (**a**).

**Extended Data Fig. 3.**
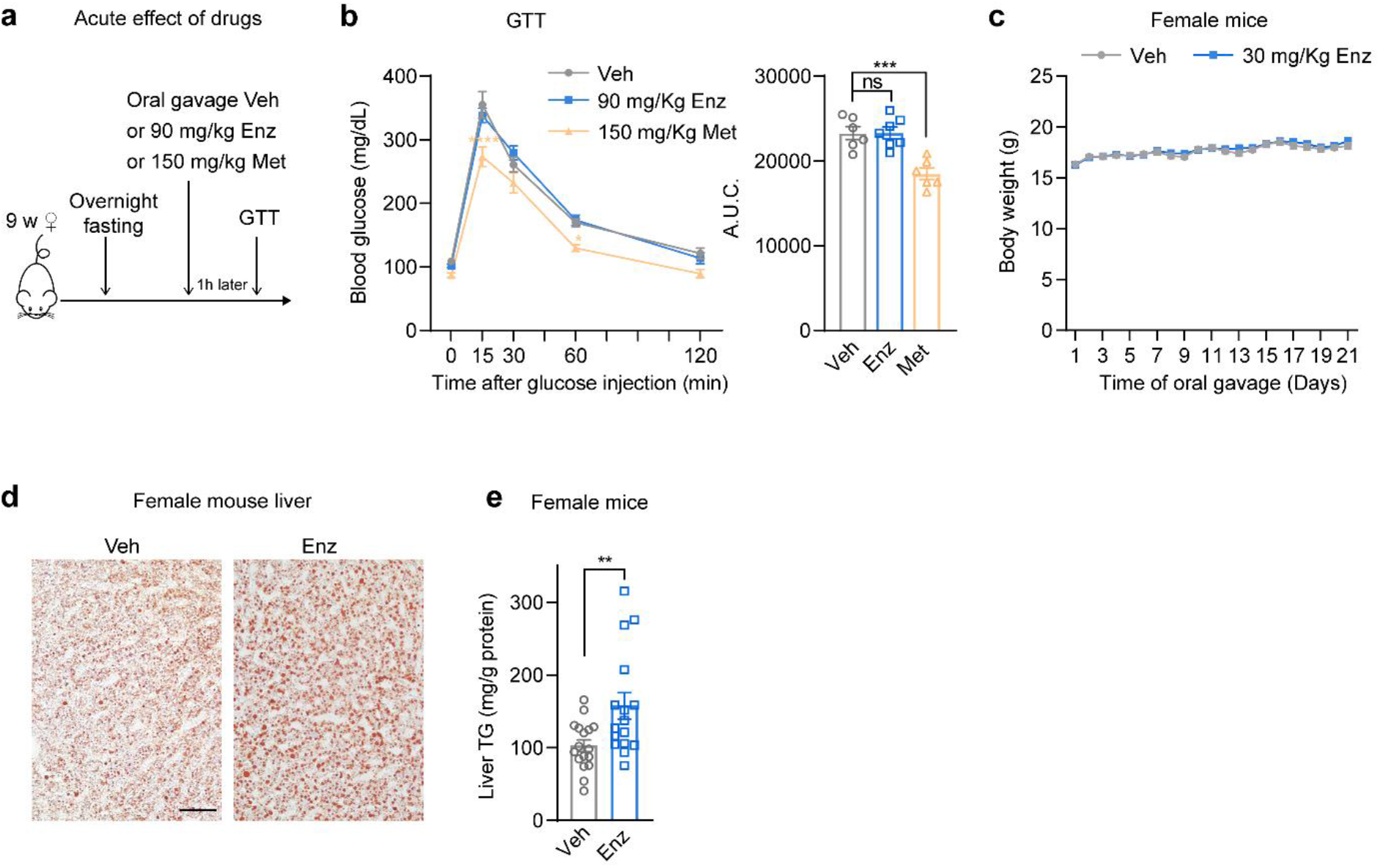
Compound test in female mice. **a,** A schematic diagram for acute effect assessments of Enz and Met on glucose tolerance test (GTT) in female mice. **b,** Blood glucose levels were measured at specified time points in female mice that were administered with Veh, Enz, or Met at indicated dosages via oral gavage, one hour prior to the intraperitoneal injection of 1.5 g/kg glucose (n=6-7 per group). The area under curve (A.U.C.) was displayed alongside the glucose curve. **c,** The body weight of female mice administered with Veh (n=9) or 30 mg/kg of Enz (n=9) via oral gavage for a period of 21 days was recorded. **d, e**, Liver fat content was evaluated in female mice treated with either Veh or 30 mg/kg of enzalutamide for a period of 3 weeks. Representative Oil Red O staining images are presented (**d**). Scale bars, 50 μm. Liver triglyceride (TG) content was quantified and expressed as a measurement (**e**). The experimental groups consisted of 16-17 mice each. The data were presented as mean ± SEM from mice with the indicated number. The statistical significance was determined using the following criteria: ***p≤0.001, **p≤0.01; ns denotes no significance with p > 0.05. The statistical tests employed were Student’s test for (**e**) and one-way ANOVA for (**b**, bar graph), while two-way ANOVA was used for (**b**, glucose curve).

**Extended Data Fig. 4.**
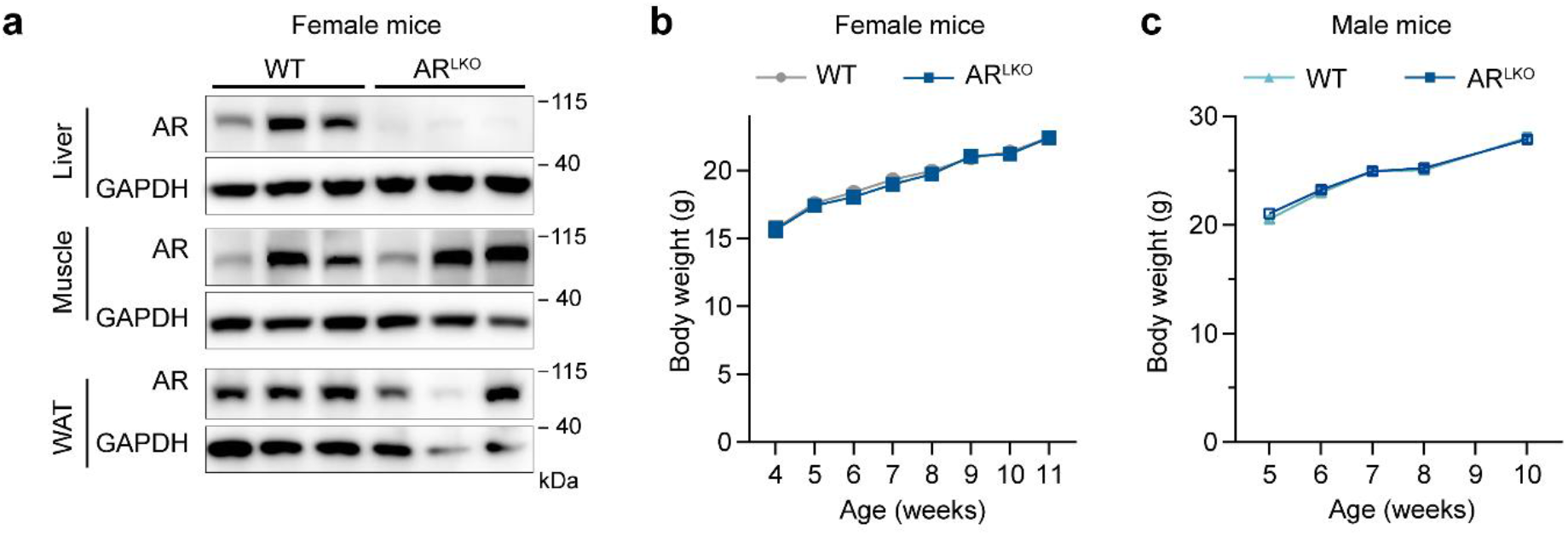
Basic characterization of the AR^LKO^ mice. **a,** The liver-specific AR knockout was confirmed through immunoblot analysis of AR protein levels in the liver, muscle, and white adipose tissue (WAT) of both WT and AR^LKO^ mice. The image presented illustrates the representative samples. **b, c,** The body weight of female (**b**) or male (**c**) AR^LKO^ mice and their WT littermates over time.

**Extended Data Fig. 5.**
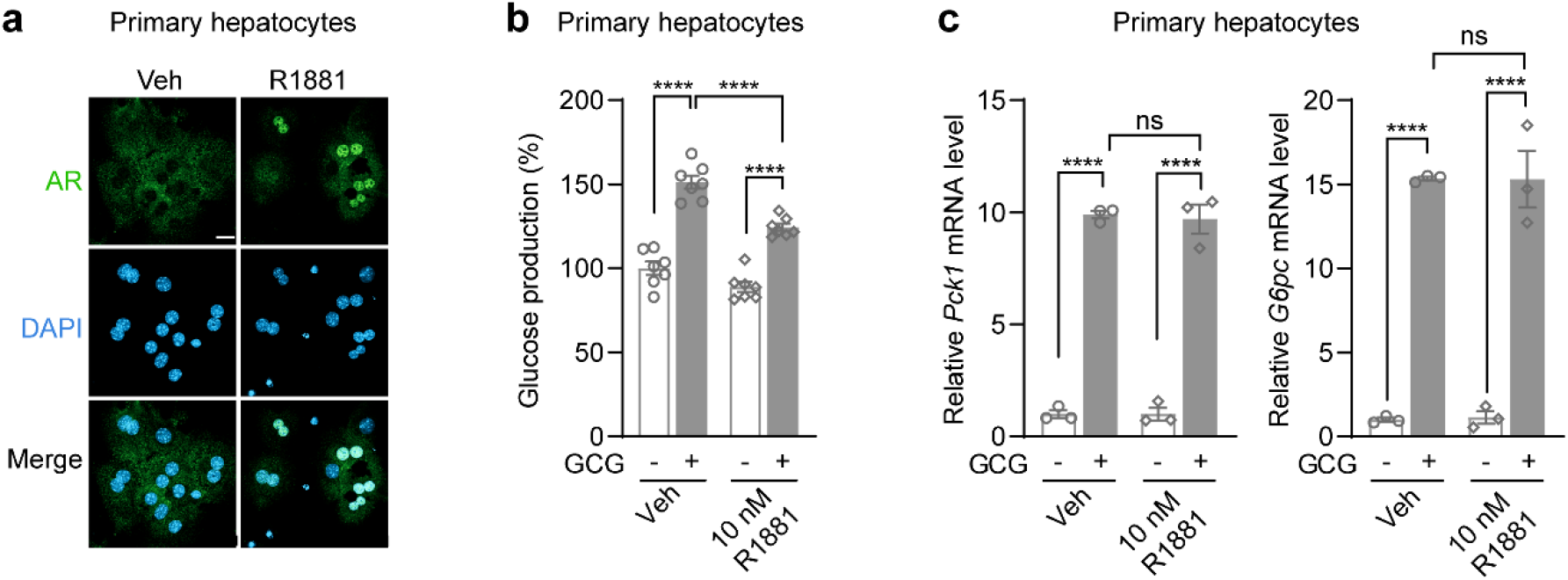
The effects of androgen on AR translocation and glucagon-stimulated gluconeogenesis in primary hepatocytes. **a,** Androgen-stimulated AR translocation from cytosol to nucleus was observed through immunofluorescence analysis of AR in primary hepatocytes of female mice. These hepatocytes were treated with either Veh or 10 nM R1881 for 24 hours. The image shown represents a representative example of the observed phenomenon. Scale bars, 20 μm. **b,c**, Hepatic glucose production (**b**) and qRT-PCR analysis of the expression of gluconeogenic genes *Pck1* and *G6pc* (**c**) in female mouse primary hepatocytes pretreated with Veh, 10 nM R1881 4 hours prior to the treatment with vehicle or 20 nM glucagon for another 18 hours. The data were presented as mean ± SEM from three or more independent experiments. The statistical significance was determined using the following criteria: ****p≤0.0001; ns denotes no significance with p > 0.05. The statistical tests employed were two-way ANOVA for (**b,c**).

**Extended Data Fig. 6.**
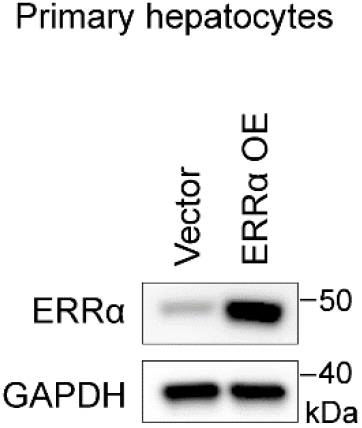
Validation of the overexpression of ERRα in primary hepatocytes. The overexpression of ERRα was confirmed through immunoblot analysis of ERRα in female mouse hepatocytes infected with Ad-Vector or Ad-Esrra.

**Extended Data Fig. 7.**
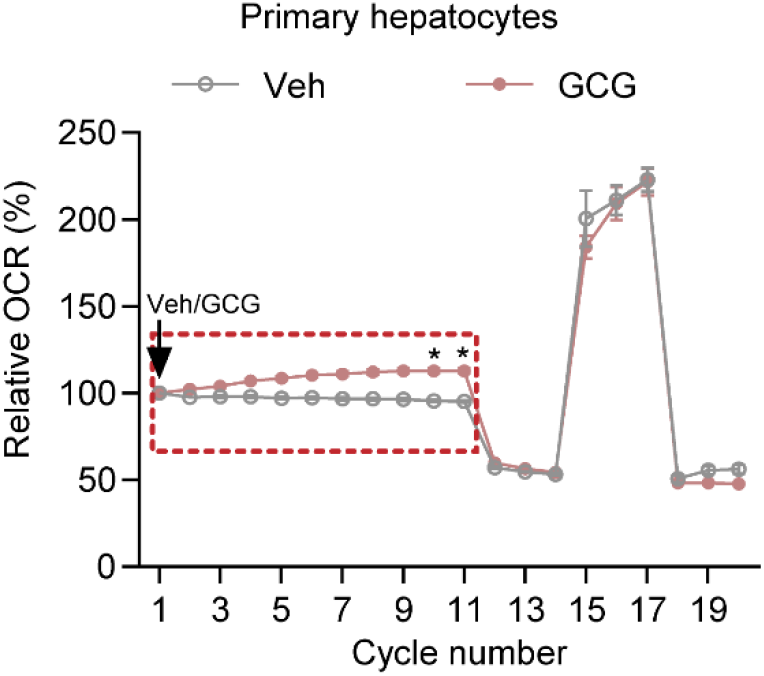
Glucagon increases the basal mitochondrial respiration of primary hepatocytes. The mitochondrial respiration of hepatocytes was measured by the Seahorse assay. During the assay, either Veh or 200 nM glucagon was injected from the beginning until the 11th cycle. Subsequently, injections of 2 μM oligomycin, 2 μM FCCP, and 1 μM rotenone/1 μM antimycin were administered for 3 cycles for each type of injection. The oxygen consumption rate (OCR) was then normalized to the corresponding 1st cycle for each group, with a normalization factor set at 100%. The data were presented as mean ± SEM from three independent experiments. The statistical significance was determined using the following criteria: *p≤0.05. The statistical test employed was two-way ANOVA.

## Supplemental tables

**Supplementary Table 3.**
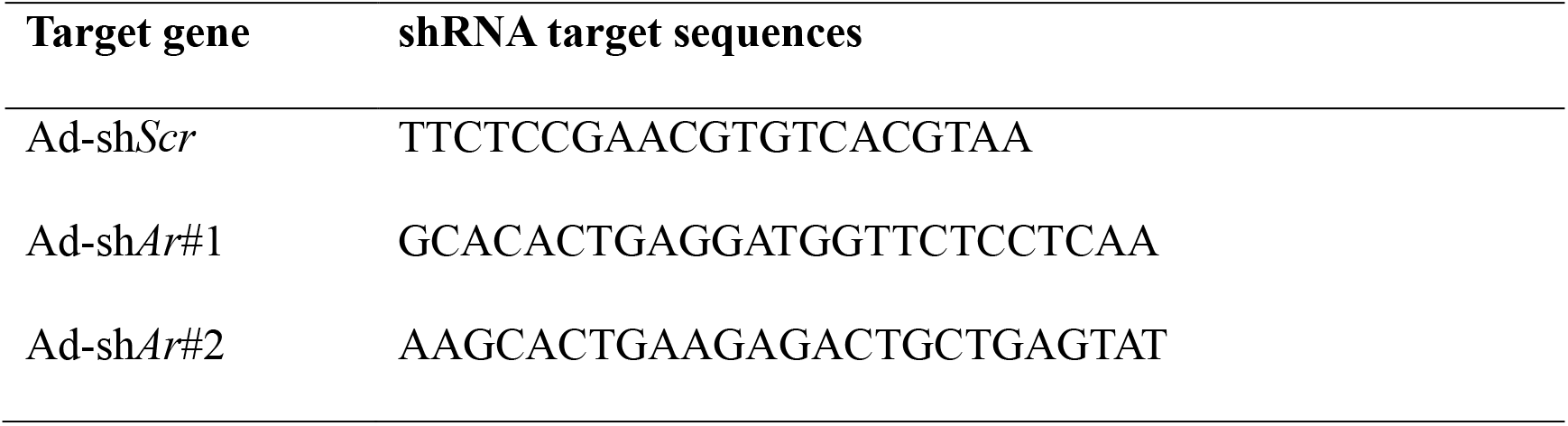
Sequences of employed shRNA.

**Supplementary Table 4.**
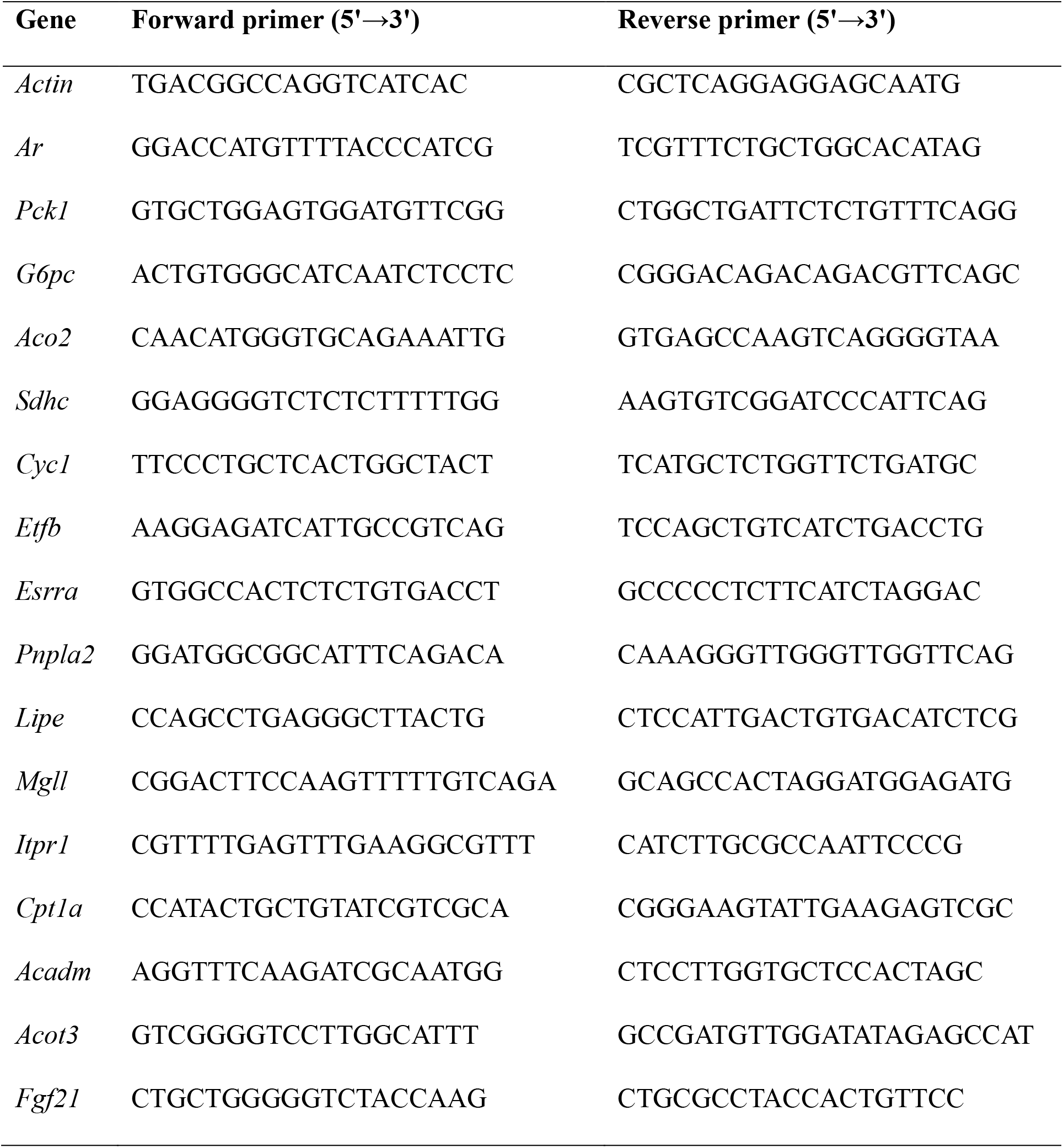
Primer sequences for qPCR.

